# Profiling crystal engineered ligands for targeting treatment resistant androgen receptors

**DOI:** 10.64898/2026.05.01.721995

**Authors:** Avan Colah, Charles Ezekiel, Sára Ferková, Pierre-Luc Boudreault, Leonard MacGillivray, William Ricke

## Abstract

Prostate cancer (PCa) is one of the principal contributors to health burden in the aging male population. PCa develops through dysregulation of androgen receptor (AR) signaling pathways. Despite improvements in diagnostic techniques and interventions, no pharmacological measures with long term efficacy have been established once PCa advances to castration resistant prostate cancer (CRPC). To circumvent this issue, tetra-aryl cyclobutanes (CBs) have been proposed as structurally distinct compounds with a mechanism of action differing from traditional androgen receptor signaling inhibitor (ARSIs). Here, we apply principles of crystal engineering and solid state synthesis to expand the class of CBs through strategic derivatization. The synthesis of the CB occurs quantitatively, producing no side products and eliminating the need for product purification. We demonstrate how head-to-tail stacking interactions of halo-pyrimidine rings can be exploited to stack and align unsymmetrical alkenes to undergo [2+2] photodimerization to generate the CB in the solid state. We examine the structure-function relationships of CBs *in vitro* by profiling AR mediated transcriptional activity, receptor translocation, and cell viability. Moreover, we explore and identify putative binding interactions within CB/AR complexes and establish an adaptive ligand-binding potential using molecular docking platforms. In total, our data suggests that CBs have unexploited therapeutic potential in CRPC and that green chemistry and crystal engineering principles offer a unique route to generating these drug candidates.

## Introduction

Prostate cancer (PCa) is the second most frequently diagnosed malignancy in men worldwide and is primarily driven by dysregulation to the androgen receptor signaling pathway (1). As PCa progresses to castration resistant prostate cancer (CRPC), most cases remain AR positive despite mechanistic deviation in AR signaling. Sequencing of advanced stage prostate cancers revealed that 20% of genomic alterations to AR were attributed to AR point mutation (2). These point mutations to the AR ligand binding domain (LBD) arise in response to the selective pressure of androgen receptor signaling inhibitor (ARSI) therapy, and result in increased receptor promiscuity and a shift in recognition of ARSIs as AR agonists rather than AR antagonists (3, 4). Examples of therapeutic shift have been reported post treatment with both first- and second-generation ARSIs including the hydroxyflutamide (Flut) induced T878A mutation (4), the bicalutamide induced W742C mutation (5), and the enzalutamide (Enza) and apalutamide induced F877L mutation (6–8). Due to the emergence of the F877L point mutation and increase in Enza resistant CRPC, darolutamide (Daro) was developed to inhibit AR LBDs with the F877L point mutation (9). Within patient populations, the prevalence of co-occurrent point mutations to AR is increasing in response to consecutive rounds of treatment with ARSIs after initial treatment failure (10). Although new small molecules like Daro can be rationally designed for clinical use, the vast majority of AR LBD targeting therapeutic bind to the LBD with a highly similar mechanism, making them susceptible to resistance point mutations (11–14). Moreover, the increased potency of second-generation ARSIs favors the emergence of resistance mechanisms since there is strong selective pressure for cell survival. Therefore, there is a critical need for a broad-spectrum AR antagonist that can inhibit a range of AR mutants consistently and with long term efficacy, now and into the future.

Recent reports have documented cyclobutane (CB) molecules as prospective drug candidates (15). CBs are experiencing an upsurge of interest in a broad range of areas related to synthetic chemistry and materials science (16). CBs are naturally formed through UV-induced intermolecular [2+2] photocycloadditions (17). The central CB ring acts as a core to emanate pendant substituents, such as aryl groups, rigidly situated in space (18). Whereas solution phase approaches to CB ring formation typically result in CB mixtures, the use of crystal engineering principals to direct formations of CBs has become highly desirable to improve product formation. CBs can now often be generated in the crystalline state as single products and in 100% yield, thus eliminating the need for compound purification (19). Moreover, the use of the solid state to form CBs of therapeutic value can be considered an unexpected, unexplored, and attractive avenue for pharmaceutical development (19). Generating cyclobutane photoproducts as AR antagonists is an exciting prospect given the remarkable ease of the solid state reactivity and the potential to form CBs that are inaccessible in solution-based approaches. This is particularly timely, as sustainable methods for generating structurally distinct therapeutic candidates are increasingly important when successfully bringing drug candidates to market.

The current understanding of CBs as point mutated AR antagonists in CRPC can be attributed to a report exploring a tetra-aryl cyclobutane scaffold as a base for drug design (20). From this work, we chose the CB scaffold 10 with -Cl substituents — which we define as the **ClCB** — as the lead compound with potential in CRPC (20). The rigidity and unique structure of **ClCB** confers a distinct mechanism of AR inhibition, whereby AR/CB complexes are retained in the cytoplasm and maintain an unliganded-like receptor conformation (20). Despite promising capacities, CBs have yet to progress to clinical trials. Therefore, complementary work building upon these initial mechanistic findings is necessary to foster development of CBs as therapeutic candidates in CRPC. Our primary goal is to better understand the potential of the **ClCB** and other halogenated CBs including the newly developed bromide -Br derivatives (**BrCB)** as drug candidates in CRPC and further establish the CB scaffold as a foundational structure for rational drug design. We demonstrate scientific advancement through our application of solid state synthesis to compound derivatization, and through the use of structural modeling to evaluate putative binding interactions between AR/CB complexes. Moreover, we define adaptive ligand binding as potential mechanistic rationale for the mutational resilience of CBs observed in cellular models of CRPC.

## Results

### Synthesis and crystal engineering of CBs

To expand our exploration of tetra-aryl cyclobutanes as candidate AR inhibitors, we synthesized the -Br styrylpyrimidine (**BrSP**) (Scheme 1, Fig. S1-S3). For an alkene to react in the solid state to form a CB, the carbon-carbon double (C=C) bonds are expected to be organized within 4.2 Å and aligned parallel (Fig 1A, S4A). We made use of the idea of steric replacement in crystal engineering to pursue the synthesis of BrSP, which would reach in the solid state to form BrCB (Table S1). Crystalline **BrSP** was determined to be photoreactive to generate the corresponding head-to-tail *rctt* isomer **BrCB** (Fig. 1E) as the solid product (Scheme 1). The crystal structures of **Cl-** and **BrSP** are isostructural. The work also describes the X-ray crystal structure of the previously synthesized **ClCB** (Fig. S4E). In the solid state, **BrSP** forms one-dimensional (1D) assemblies held together by C-H…N hydrogen bonds involving the pyrimidine moieties (Fig 1B, 1C, S4B, S4C). X…X halogen bonds effectively align the 1D assemblies such that the C=C bonds are separated by 3.37Å and fall within the geometry criteria for a [2+2] photodimerization (4.2Å and parallel) (Fig. 1D, S4D). UV-irradiation of **BrSP** results in quantitative conversion of the solid to **BrCB** (Fig. S3). The CB in the solid state was determined to interact with its neighbor through C-H…N hydrogen bonds to form planar sheet (Fig. 1F, S4F). We emphasize the utility of generating and profiling the **BrCB** as the compound also offers distinct potential both as a standalone drug candidate and as a base scaffold for future chemical optimization (e.g. Suzuki coupling) (21).

**Scheme 1.**
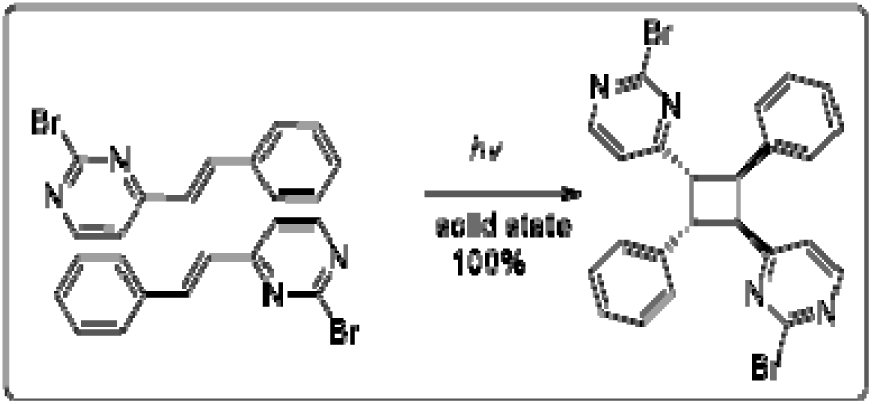
2+2 photodimerization of BrSP to generate **BrCB** in the solid state.

**Figure 1.**
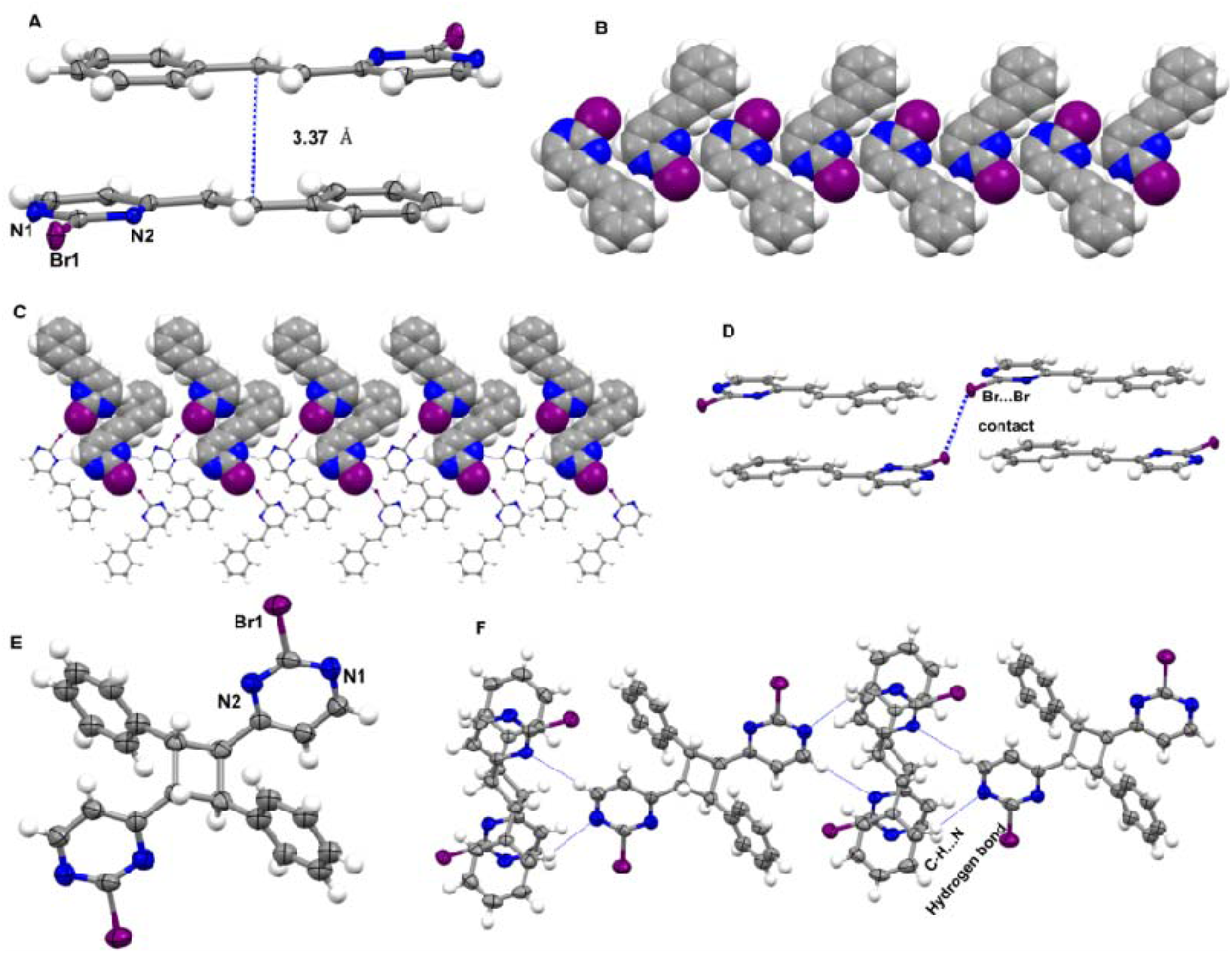
Crystal structures BrSP and **BrCB**. **(A)** head-to-tail dimer showing the stacking of the C=C bonds of the alkene with a 3.37 Å distance **(B)** 1D edge-to-edge view of sheets encompassing neighboring molecules of BrSP supported by C-H‧‧‧ N hydrogen bonds **(C)** 2D C-H‧‧‧ N and Br‧‧ ‧ Br intermolecular interactions **(D)** Br‧‧‧ Br halogen bond of the alkene of the nearest neighbor BrSP **(E)** ring of **BrCB** highlighting the rctt stereochemistry, and **(F)** extended packing of the **BrCB** showing the interaction of the nearest neighbor through two C-H‧‧‧ N hydrogen bonds. Corresponding color identifiers for atoms include hydrogen (white), carbon (grey), nitrogen (blue) and bromine (purple). All intermolecular interactions are indicated by dotted blue lines.

### The ClCB and derivative BrCB exhibit potent dose dependent transcriptional inhibition

Previous **ClCB** (CB 10) studies were conducted using AR transfection models and a mixture of PCa cell lines (20). However, much of this work utilized models where point mutated ARs were not expressed endogenously. Our goal is to prioritize consistent model systems where AR and downstream cellular machinery will be most representative of the CRPC cellular environment. To examine each ligand-receptor complex *in vitro*, we employed CRPC cell lines that endogenously express AR point mutants. The LNCaP (AR^T878A^) was chosen to represent single point mutation (22, 23). The MR49F (AR^F877L/T878A^) cell line — a derivative of LNCaP—was used to represent broader drug resistance to two ARSIs (24). This is the first report for the application of CBs to CRPC models with co-occurrent AR LBD point mutation. For AR^WT^, an LNCaP derivative cell line (LNCaP/AR) was used where a codon switch was induced to express AR^WT^ (25). Functional validation confirmed that LNCaP/AR cells were susceptible to inhibition by flutamide due to loss of T878A point mutation, and enzalutamide was demonstrated to induce transcriptional activity in the MR49F cell model where AR^F877L/T878A^ is present (Fig. S5).

Structural comparison of the AR LBDs present in our cell lines of interest showed that the LBD of the wild-type (WT) AR, located at its *C-terminal* region, comprises 11 α-helices, 4 β-sheets, and 14 turns. The T878A mutant exhibits a structural rearrangement resulting in the presence of 12 α-helices and 15 turns, while the F877L/T878A double mutant displays 16 turns. Structural superposition of the WT (blue), T877A (cyan), and F877L/T878A (purple) AR LBDs reveals conformational deviations localized within the disordered loop region (Fig. 2A). The structural deviations highlight the conformational flexibility of the AR LBD induced by point mutations, which may influence ligand binding and receptor activation.

**Figure 2.**
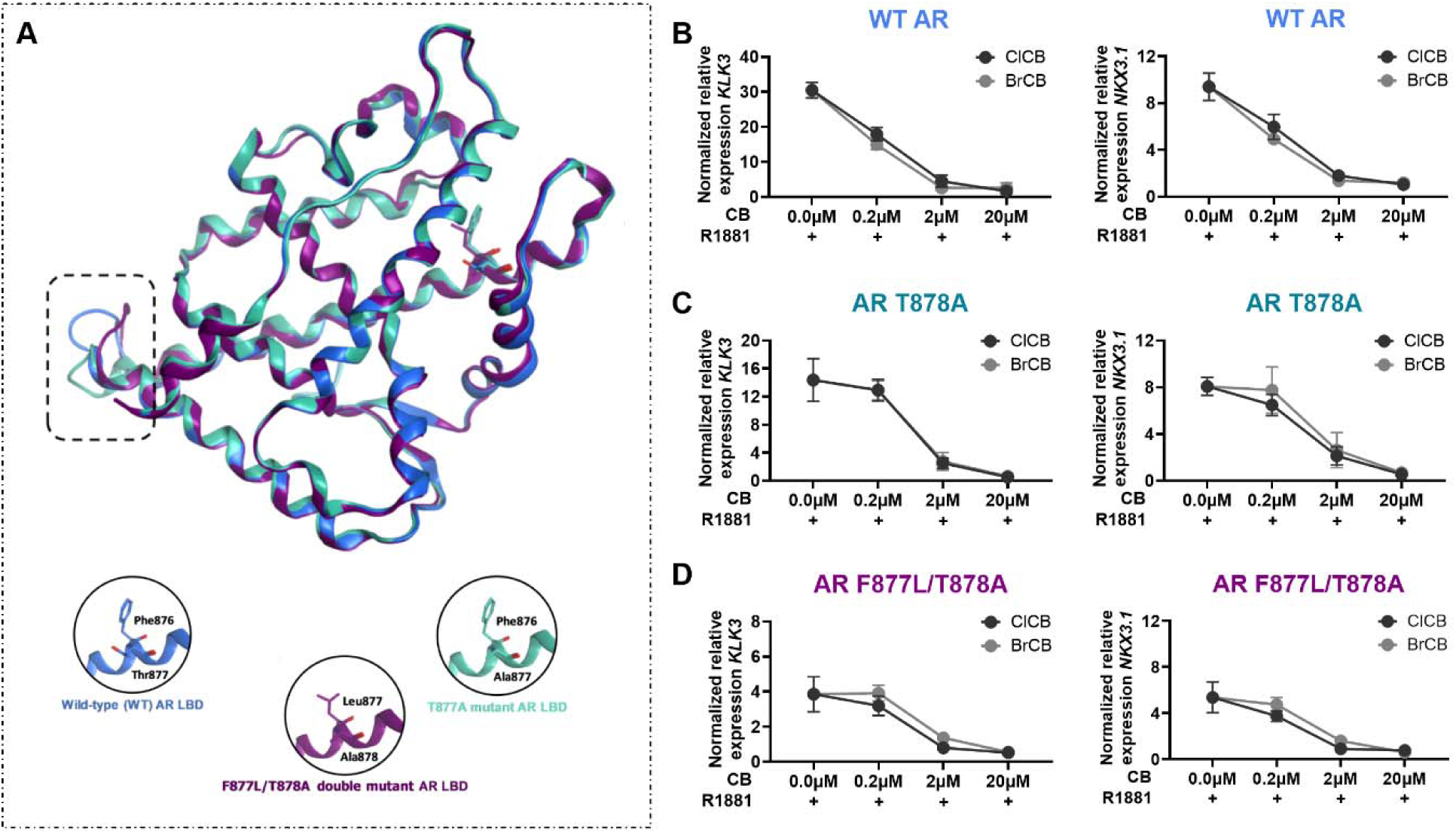
Dose dependent transcriptional inhibition by CBs. **(A)** structural overlay of the WT (blue), T877A mutant (cyan), and F876L/T877A double mutant (purple) AR LBDs. Disordered overlapping regions are highlighted in a black square. The three LBDs are representative of the structural changes to AR in the LNCaP/AR, LNCaP, and MR49F cell lines. **(B-D)** qPCR quantification of KLK3 or NKX3.1 transcript expression after 24hr treatment with 0.2µM, 2µM, and 20µM ClCB or BrCB in the presence of 1nM R1881. Observations are reported for LNCaP/AR (ARWT), LNCaP (ART878A) cells, and MR49F (ARF877L/T878A) cell lines respectively.

Hallmarks of AR activation include transcription of *KLK3* and *NKX3.1* target genes. Quantification of gene expression determined the dose of **ClCB** and **BrCB** necessary for complete inhibition of AR regulated transcription in each cell line. In all three cell lines, dose dependent inhibition of AR target gene expression was observed for the **ClCB** and **BrCB** (Fig 2B-D).

### CBs function to inhibit AR nuclear internalization and transcriptional activity

The LNCaP (AR^T878A^) and MR49F (AR^F877L/T878A^) models were chosen to evaluate CBs influence on cellular processes downstream of therapeutic resistant AR. Importantly, both the LNCaP and MR49F cellular models endogenously express their AR point mutants and exhibit AR amplification which often co-occurs clinically along with point mutation (10). These models will provide the most representative *in vitro* systems for evaluating downstream molecular mechanisms.

One of the primary methods anti-androgens inhibit AR is through a blockade of nuclear internalization (25, 26). As reported for the **ClCB** (20), we hypothesized that blockade of AR nuclear translocation would also occur with the **BrCB** owing to the structural similarity of the two compounds. LNCaP and MR49F cells were stained via fluorescence immunocytochemistry for AR and counterstained for nuclei with DAPI. Confocal imaging and analysis of LNCaP and MR49F cells showed that AR was ∼35% cytoplasmic and ∼65% nuclear at baseline in both models. Treatment with R1881 stimulated nuclear internalization of ARs, and cotreatment of R1881+CB or ARSI halted this effect in both cell lines (Fig. 3A). Quantification of cell compartmentalization by colocalization of ARs to DAPI was measured with the Mander’s 2 coefficient (Fig. 3B) and the nuclear/cytoplasmic fraction of ARs (Fig. 3C). Together, these findings indicate that CBs block AR nuclear internalization and that progression from isolated to co-occurrent point mutation does not hinder the CB mechanism of action.

**Figure 3.**
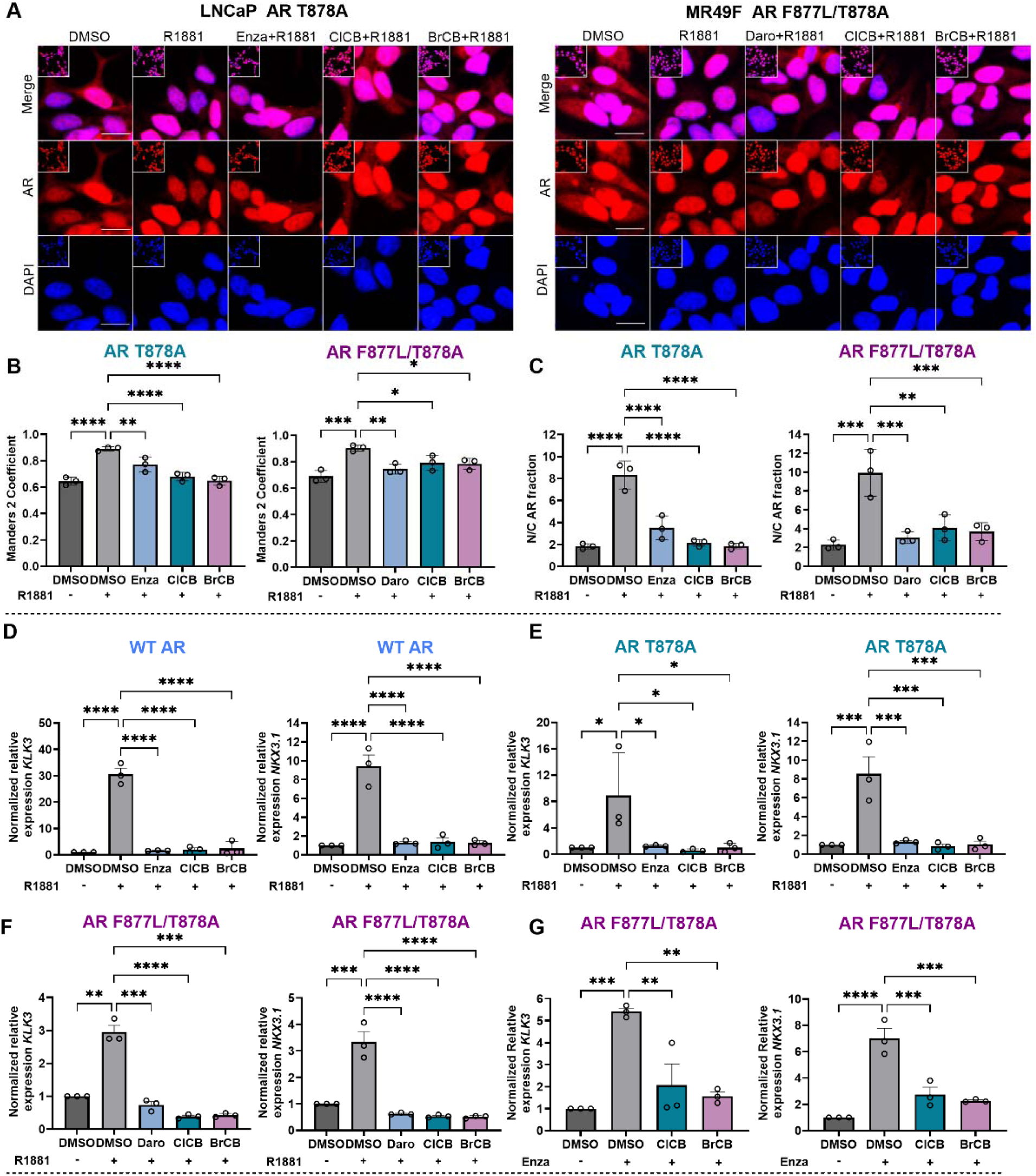
BrCB. mechanism of action is consistent with **ClCB** and analogous to the impacts of traditional ARSIs. **(A)** Representative confocal ICC images of LNCaP (ART878A) cells and MR49F (ARF877L/T878A) after 4hr treatment with 20 µM CBs or ARSIs. AR protein (red) and DAPI as a counterstain for nuclear area (blue). Scale bar = 20µm. **(B)** Quantification of AR protein colocalized with DAPI represented with the Mander’s 2 coefficient as the proportion of nuclear AR in LNCaP and MR49F cells. Treatment with CBs blocks nuclear internalization of AR induced by R1881. **(C)** Secondary validation of cellular AR protein localization assessed using the nuclear/cytoplasmic fraction (N/C fraction) of AR staining. (D-F) KLK3 and NKX3.1 gene expression demonstrating blockade of the AR signaling pathway after 24hr combination treatment with 20µM ClCB or 20µM BrCB +1nM R1881 compared to the FDA approved ARSI 20µM + 1nM R1881 in the LNCaP/AR (ARWT), LNCaP (ART877A) cells, and MR49F (ARF877L/T878A) cell lines. (G) Quantification of gene expression downstream of AR in MR49F ARF877L/T878A cells where transcription was induced with 20µM Enza and subsequently blocked by cotreatment of 20µM CBs + 20uM Enza.

Independent studies on each cell line were completed to gauge the inherent activity of each ligand as a full or partial agonist or antagonist. Representative images and quantification of AR localization in each cell line was indicative of complete antagonistic properties for both the **ClCB** and **BrCB** (Fig. S6, S7). Consistent with expectations, R1881 and Flut promoted nuclear internalization of AR in LNCaP cells while R1881, Flut, and Enza increased nuclear AR in MR49F cells. All other ligands did not induce substantial changes in AR cell compartmentalization. (Fig. S6,S7).

Our work compared CBs at matched concentrations of 20 µM to ARSIs. Enza was used as the control ARSI in LNCaP and LNCaP/AR cell lines, and Daro was used as the control ARSI for MR49F cells. Consistent with expectations, treatment with R1881 induced the expression of *KLK3* and *NKX3.1* and co-treatment of R1881+CBs blocked transcript expression with similar efficacy to traditional ARSIs (Fig. 3D-F). Upon exposing the Enza resistant MR49F cell line to 20 µM Enza, AR signaling was activated, and CBs were demonstrated to outcompete Enza and block transcription downstream of AR (Fig. 3G). These findings highlight CBs as drug candidates with comparable inhibitory efficacy to second generation ARSIs.

### CBs exhibit characteristics of cytostatic rather than cytotoxic agents

A characteristic impact of AR inhibition in AR dependent cells is loss of cell viability. We expected that dose dependent inhibition of AR target gene transcription would render a similar trend in cell viability when measured by MTT assay. LNCaP/AR, LNCaP, and MR49F cells were treated with 0.01 - 100µM **ClCB**, **BrCB**, or Daro to evaluate cell viability after 96 hr of treatment (Fig. 4A). Surprisingly, cells treated with CBs retained their viability up to the maximum solubility of CBs at 30µM in media. Matched concentrations of Daro produced the expected dose response curve and reduced viability with increasing concentration in all three cell lines. These data demonstrate that CBs exert distinct impacts on cell viability when compared to traditional ARSIs despite potent AR transcriptional inhibition. This finding points to potential cytostatic rather than cytotoxic activity of CBs. To further evaluate this hypothesis, we examined the cell cycle markers *CDC20* and *UBE2C* since cytostatic agents prevent cancer progression through inhibiting cell growth and division rather than inducing cell death. Quantification of *CDC20* gene expression in the LNCaP and MR49F cell lines after CB treatment demonstrated a reduction in the mRNA encoding *CDC20*, an essential regulator of cell division (anaphase initiation), compared to active cycling induced by R1881 (Fig. 4B). The cell cycle regulator *UBE2C* was also demonstrated to decrease compared to R1881 after CB treatment in the LNCaP cell line, but not the MR49F cell line (Fig. 4C). Together, this data supports a cytostatic functionality of CBs where cell growth is reduced but cell viability is retained compared to cytotoxic agents like ARSIs.

**Figure 4.**
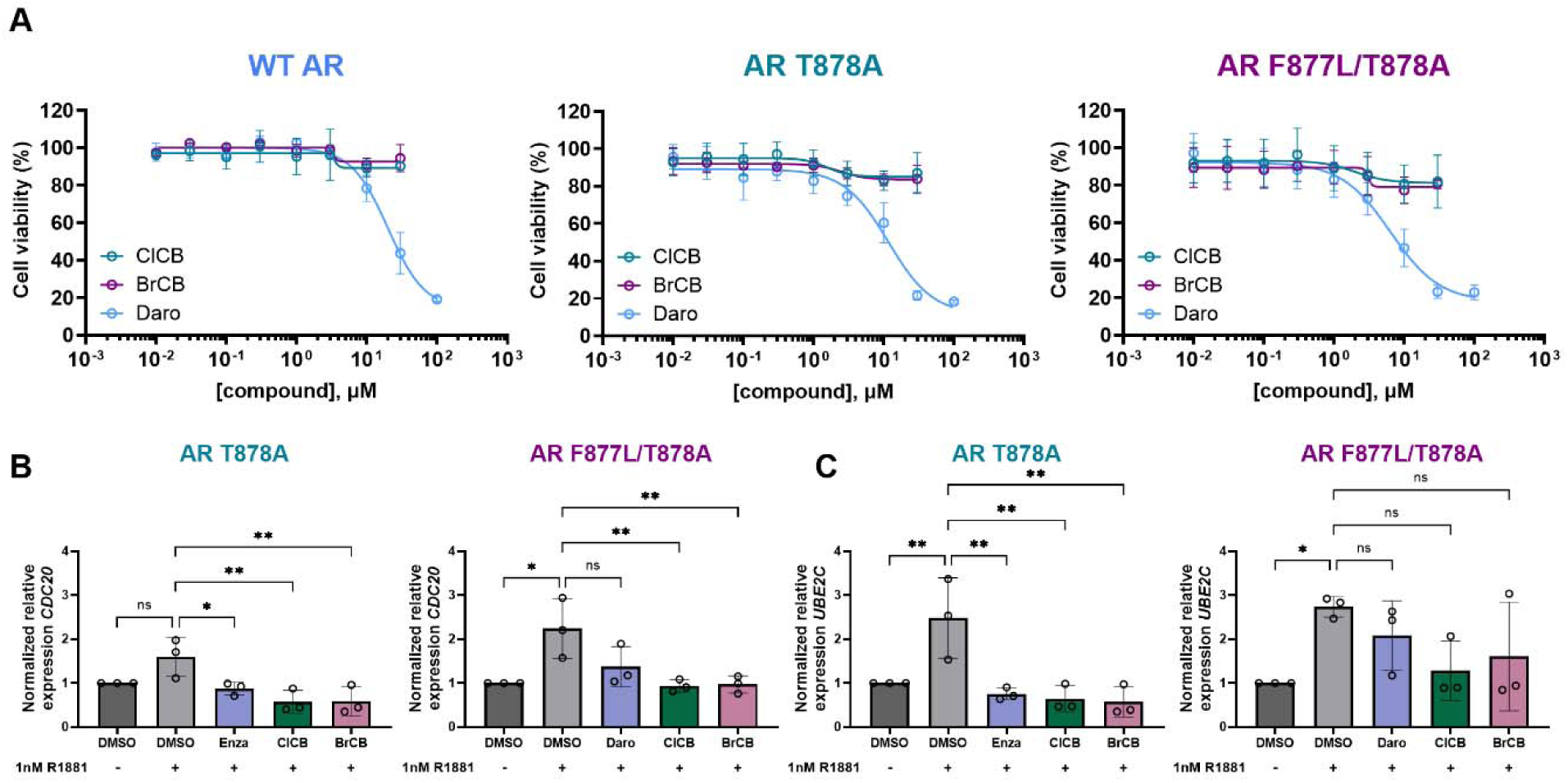
CBs exhibit characteristics of cytostatic agents **(A)** MTT assay data evaluating percent cell viability in all cell lines after 96hr of exposure to increasing concentrations of CBs or Daro. A dose response curve is observed with Daro treatment but not with CB treatment. **(B)** Quantification of *CDC20* gene expression in the LNCaP and MR49F demonstrating a reduction in cell division compared to R1881 after 24hr treatment (C) Expression of the cell cycle regulator *UBE2C* was decreased compared to R1881 after 24hr treatment in the LNCaP but not the MR49F cell line.

### Molecular docking and comparison of CB binding poses within AR LBD point mutants

As CBs exhibit a distinct structure and cellular impact from both first- and second-generation ARSIs, we sought to define CB/AR ligand-receptor complexes. As reported, resistance-associated point mutations within the AR LBD—notably T878A and F877L—can confer therapeutic resistance by altering ligand recognition or modulating receptor activation. In some cases, resulting in a functional switch from antagonist to agonist behavior (12–14). In the present study, molecular docking was carried out to elucidate the binding modes of **ClCB** and **BrCB** ligands within the wild-type (WT) AR and two mutants, and to systematically evaluate the impact of progressive resistance mutations on binding of both ligands. Literature on AR point mutations can differ in annotation based on codon counts used. Therefore, Phe can be located at either the 876 or 877 codon and Ala can occur at either the 877 or 878 codon. F876L and F877L are equivalent point mutations, as are T877A and T878A. Our molecular docking figures employ codon counts consistent with the PDB files used for analysis.

Hydrophobic contacts, aromatic interactions and hydrogen bonding (H-bond) networks established between the ligands (**ClCB** and **BrCB**) and ARs were analyzed in depth using the “Protein Contacts” function in MOE software to give, with three-dimensional (3D) representations (Fig. 5) and complementary two-dimensional (2D) ligand interaction diagrams (Fig. S8). Both ligands were found to occupy the lipophilic pocket of ARs, predominantly formed by nonpolar amino acid residues, (i.e Leu, Phe, Trp and Val) (Fig. 5, S8). This spatial arrangement suggests that weak, non-directional Van der Waals (VdW) forces constitute the principal energetic contributors to ligand–receptor complex formation. Given the pronounced ligand hydrophobicity, the binding process is primarily entropically driven, thereby conferring a thermodynamically favorable interaction profile. Nonetheless, in some cases, additional stabilizing, directional interactions—such as H-bonds and aromatic interactions—were also determined upon docking (Fig. 5, S8). For example, in the AR^WT^, the Cl-pyrimidyl moiety of **ClCB** participates in weak H-bonding interactions with Met742 (d_H-SD_ = 3.00 Å) and Met787 (d_H-SD_ = 2.64 Å), each contributing to an estimated interaction energy of –0.50 kcal/mol (Fig. 5A, 6A, S8A). Structural superposition of the **ClCB** ligand in both the WT and T877A mutant AR LBDs reveals a highly conserved binding mode, characterized by the co-localization of the phenyl rings and the electron-deficient heteroaryl moieties within the binding pocket (Fig 6A, C). In contrast, within the F877L/T878A double mutant AR, the Cl-pyrimidyl group engages exclusively in a weak hydrogen bond with Met788 (–0.80 kcal/mol, d_H-SD_ = 2.46 Å), failing to establish contact with Met743 (Fig 6B, S8E, Table S2a-d).

**Figure 5.**
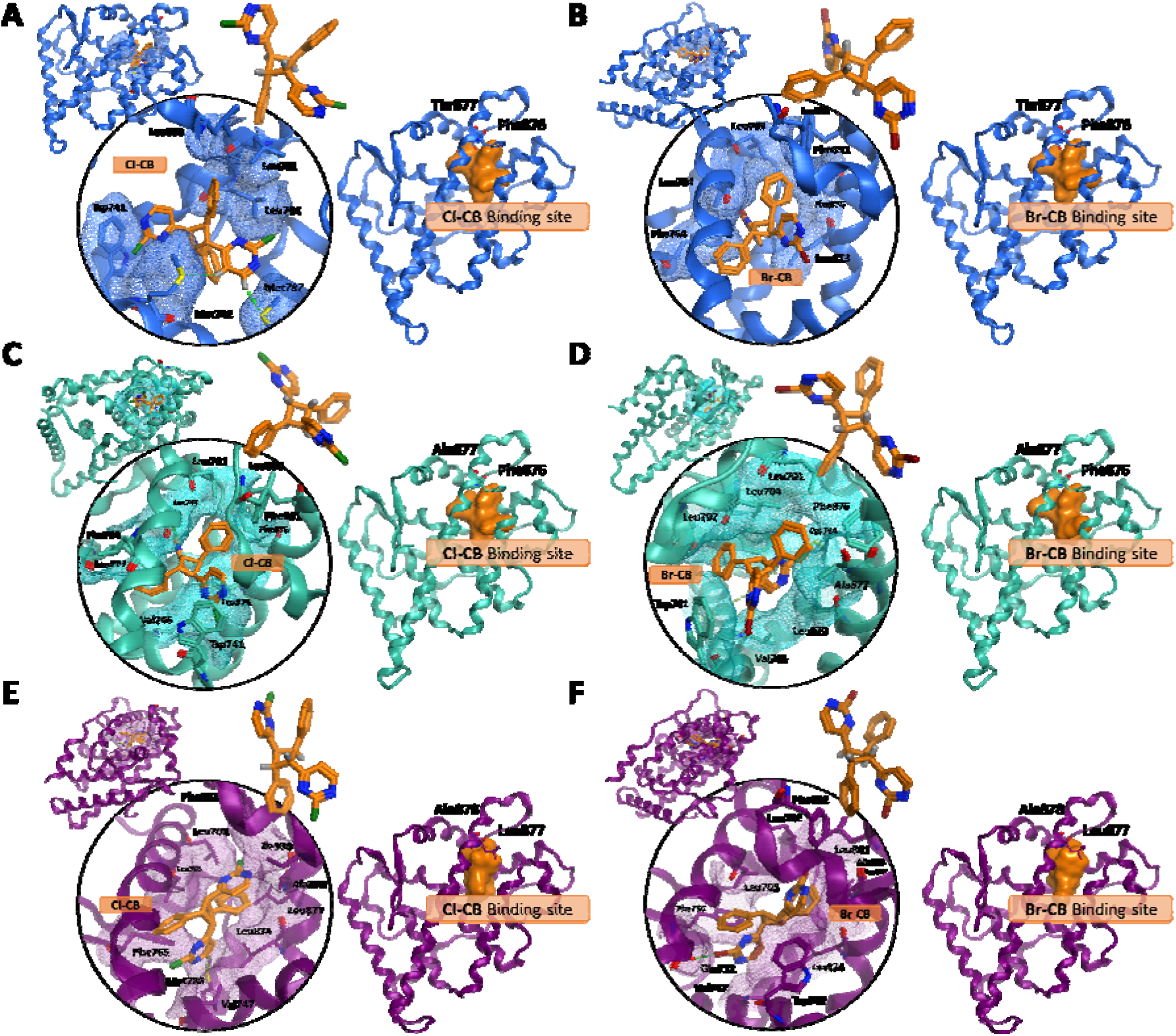
Docking poses of CBs are energetically favorable in LBDs of AR WT, ART877A, and ARF877L/T877A. Comparative analysis of ClCB and BrCB docking poses in the **(A, B)** WT (blue), **(C, D)** T877A mutant (cyan), and **(E, F)** F877L/T878A double mutant (purple) AR LBDs. Black circles highlight the detailed binding interactions of **ClCB** and **BrCB** within the lipophilic pocket of the AR LBDs. Sidechains of residues involved in hydrophobic interactions with the ligands are represented as bold sticks in blue, cyan, or purple, corresponding to each receptor variant, and are annotated accordingly. The spatial contour of the lipophilic pocket accommodating each ligand is visualized as a semi-transparent mesh. H-bonds are illustrated as green dashed lines with a strength bar, while aromatic interactions are shown as light green dashed lines. Each annotated docking pose is paired with a surface model depiction for each respective ligand providing spatial context for their interactions within the ribbon representation of the receptor. The F876L and T877A mutations are highlighted as bold sticks in the AR mutants and the corresponding WT receptor, emphasizing their structural positioning within the binding domain. Note: F876L/F877L and T877A/T878A are used interchangeably to annotate point mutations throughout the literature dependent on method of codon count.

**Figure 6.**
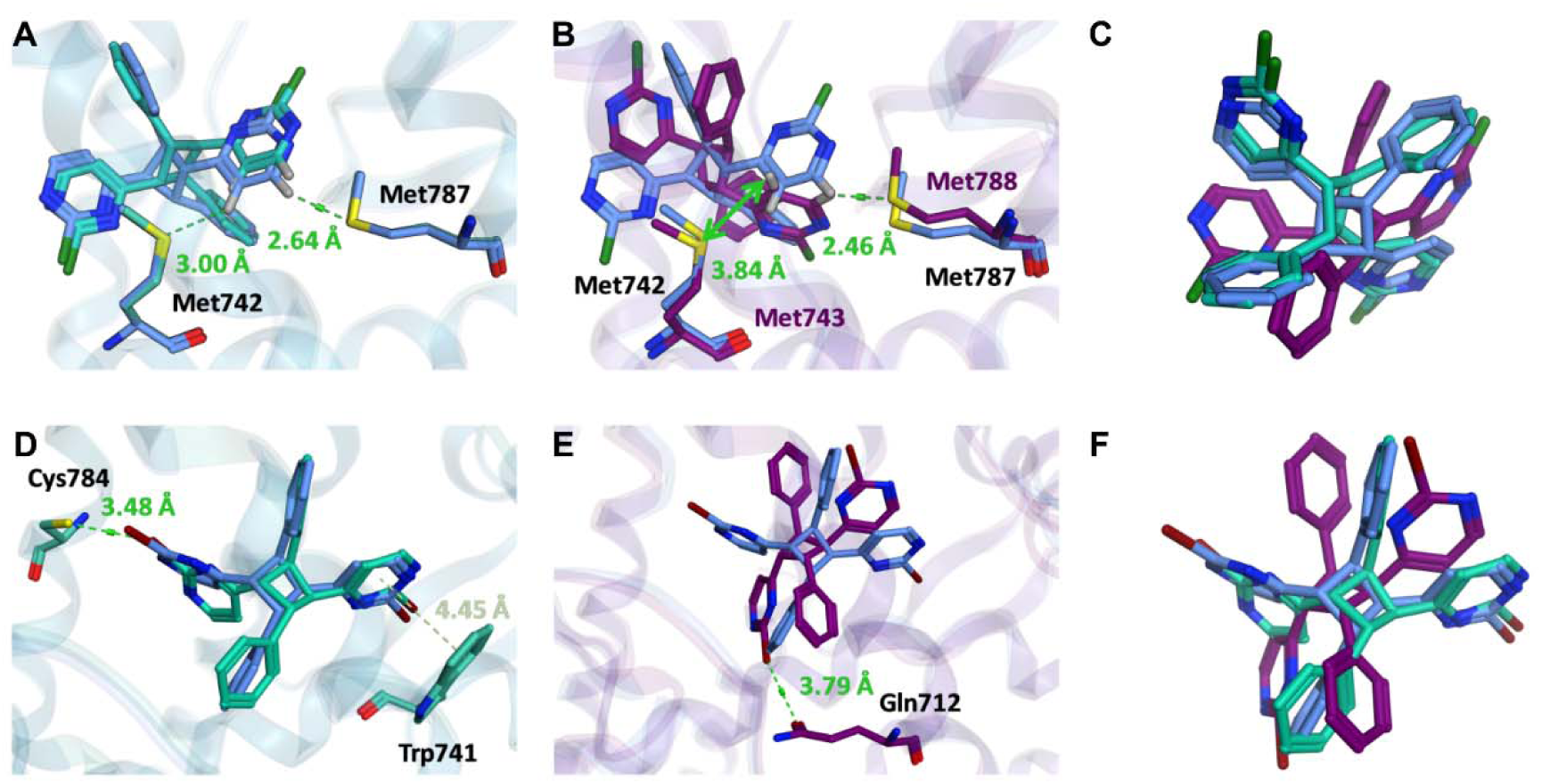
Comparative binding poses of **ClCB** and **BrCB** within AR LBD point mutants. **(A, D)** Structural superposition of **ClCB** and **BrCB** docking poses within the wild-type (WT, blue) and T877A mutant (cyan) AR LBDs. The off-set stacking aromatic interaction is indicated by a light green dashed line. **(B, E)** Superposition of **ClCB** and **BrCB** ligand docking conformations within the WT (blue) and F877L/T878A double mutant (purple) AR LBDs. The spatial distance from the hydrogen atom of the Cl-pyrimidyl moiety in the docked **ClCB** to Met743 is indicated by a green bidirectional arrow. Predicted H-bonding and VdW interactions between ligands and AR LBD variants are illustrated as green dashed lines with a strength bar, with the associated bond distances labeled in Å. The AR LBDs are rendered in transparent ribbon representations to emphasize ligand orientations across the AR LBD variants. **(C, F)** Overlay of docked **ClCB** and **BrCB** conformations across all three AR LBD variants: WT (blue), T877A mutant (cyan), and F877L/T878A double mutant (purple).

The docking poses of **ClCB** and **BrCB** in both the WT and T877A single mutant AR LBDs revealed a high degree of structural similarity, indicating that the Thr877 to Ala877 substitution exerts minimal influence on ligand orientation or binding site occupancy (Fig 6 A, B, D, E). In contrast, docking within the F877L/T878A double mutant AR LBD demonstrated a notable conformational shift in both ligands. This is attributable to altered steric constraints introduced by the combined mutations F877L/T878A, which reshape the binding cavity and induce ligand reorientation (Fig. 6 C, F). The calculations offer foundational insight into the forces driving favorable CB/AR LBD interactions across point mutants. With this computational data, informed design can guide the synthesis of **BrCB** and other halogenated derivatives with enhanced pharmaceutical potential. Strategic functionalization should continue to prioritize ligands that maintain antagonism even in the event of dramatic ligand reorientation. This unique property of CBs is an unconventional advantage that fosters a new perspective on rational drug design for CRPC.

## Discussion

Here, we describe the application of CBs in models of CRPC, involving both isolated and co-occurrent AR point mutation, to examine the ability of CBs to bypass mechanisms of therapeutic resistance. Our results led us to consider CBs as cytostatic rather than cytotoxic agents. Consistent with the expectations of cytostasis, CBs treatment induced markers of slowed growth in CRPC cells without killing them at concentrations sufficient to block AR activity. Moreover, CBs accommodate structural changes in the AR LBD through altered binding orientation as demonstrated when modeled in AR progression from isolated to co-occurrent LBD point mutation. This molecular feature is highly desirable and could offer the first step in optimization of an adaptive AR inhibitor that is less susceptible to the pharmacological impacts of point mutation. However, additional structural biology experimentation is necessary to further validate CBs as adaptive ligands. Collectively, our findings demonstrate that strategic derivatization of CBs, using crystal engineering principles, can expand this distinct class of AR inhibitors. CB scaffolds harbor unique pharmaceutical potential which should continue to be explored using rational drug design tactics. Our work endorses CBs as desirable therapeutic candidates in CRPC and other androgen driven diseases and disorders.

Crystal engineering is emerging as a technique to synthesize functional materials with applications in pharmaceutics, energy storage, agriculture, and electronics (27). Reactions in crystals are now providing access to molecules difficult to obtain using conventional (solution-based) organic synthesis. Solid state reactions help to perform organic syntheses that are efficient, sustainable, and atom-economical (28). Given that CB products are generated quantitatively, without side products, and with pharmacological activity against AR, it is evident that this is a vital development in synthesizing therapeutic agents for CRPC. Since capitalizing on thoughtful, resource efficient, and environmentally conscious synthesis is becoming a priority in academic research and industries (28), research that emphasizes solvent-reduced and solvent-free reactions is important (28). Considering CBs as potential candidates in the pharmaceutical development pipeline, the [2+2] photodimerization reaction (Scheme 1), inherently incorporates sustainable synthesis. Notably, this reaction is accomplished through UV radiation, is atom economical, and produces a pure product with no byproducts.

Thus far, application of crystal engineering to the pharmaceutical industry has generated a class of multi-component drugs known as cocrystals (29). Accumulating successes of cocrystals highlights potential for crystal engineering to impact current approaches to rational drug design (29). However, crystal engineering of reactions in the solid state that allow for the development of single component drug molecules has yet to gain similar traction. To date, publications leveraging crystal engineering techniques in pharmaceutical development for CRPC are limited. Therefore, it is significant to report that the therapeutic candidates **ClCB** and the newly derived **BrCB** are generated using crystal engineering. Moreover, abundant opportunity remains to further optimize CBs through the combination of structure activity relationship analysis and crystal engineering.

In the context of CRPC, CBs are attractive drug candidates with thermodynamically favorable interaction profiles in AR LBDs. Our X-ray crystal data have allowed for close examination of the structure activity relationships between each CB paired with AR^WT^, AR^T878A^, and AR^F877L/T878A^. Both the **ClCB** and **BrCB** take on a distinct ligand binding orientation in context of co-occurrent point mutation (Fig.5). Historically, molecular docking studies have shown that ligand reorientation of ARSIs including flutamide, bicalutamide, and enzalutamide leads to a loss of function in respective point mutated AR LBDs (11–14). The functional shift is primarily a result of the inability of ARSIs to sterically hinder helix 12 (H12) from closing after point mutation. Once closed, H12 forms a coactivator binding site necessary for transcription (11–14). Despite co-occurrent point mutation inducing reorientation of the CBs, we find that complete inhibition of AR transcriptional activity (AR target genes *KLK3* and *NKX3.1*) and blockade of AR nuclear internalization still occurs (Fig 2,3). This remarkable finding indicates that the structure of CBs offers an adaptive inhibitory profile in contexts where traditional ARSIs are prone to failure. Although our work has yet to profile H12 dynamics in the context of CB/AR complexes, we propose this area as a next critical component necessary to optimize CB scaffolds as therapeutic candidates in CRPC. Moreover, CryoEM experiments will be necessary to bolster the evidence that CBs may have adaptive ligand binding functionality.

During our profiling of the CBs in models of AR LBD point mutants, we predicted – based on the foundational biological concept that form dictates function – that **BrCB** would exhibit major functional overlaps with the **ClCB**. Comparable pharmacological profiles were observed for the two compounds. Overlapping capacities include pan-inhibition of AR LBD point mutants despite ligand reorientation, dose-dependent AR inhibition, blockade of AR transcriptional activity and AR nuclear entry, and minimal impacts on cell viability. The absence of an impact on cell viability led us to consider cytostatic rather than cytotoxic functions of CBs. Consistent with expectations of cytostatic agents, the cell cycle markers *CDC20* and *UBE2C* were reduced after CB treatment demonstrating slowed growth. This finding was consistent with previous reports demonstrating that the ClCB suppresses cellular proliferation *in vitro* (20). Cofactor profiling of AR/ClCB complexes offers a plausible explanation for retention of cell viability after ligand binding (20). The AR/ClCB ligand complex was found to be conformationally similar to an unliganded-like receptor and did not cluster with the receptor conformation induced by traditional first- or second- generation ARSIs (20). Therefore, cellular mechanisms downstream of AR cofactors that reduce viability after ARSI treatment may not be triggered when CBs bind to AR. This concept provides an interesting route for future investigation of how the structural overlap between the ClCB and BrCB guides similar cellular mechanisms while simultaneously remaining distinct from traditional ARSIs.

To highlight the pharmaceutical potential of CBs across the diverse mutational landscape of the AR LBD, we reiterate the originality of deciphering a structure that can effectively bind to the AR LBD in multiple orientations while retaining antagonistic function and potency. Moreover, our molecular docking studies provide a basis for ligand optimization from the CB scaffold. Computational data implies that binding affinity and selectivity could be significantly enhanced through strategic functionalization at the ortho/para-position of the phenyl ring of CBs with H-bond acceptor/donor groups. These conclusions highlight both immediate and continued potential for the solid state development of CB derivatives with distinct pharmaceutical potential.

Our work is particularly timely as many clinically prevalent AR LBD targeting therapeutics prioritize potency and ligand functionality in highly specific scenarios (9, 30, 31). This technique for novel therapeutic development has resulted in the cyclical failure of ARSIs since potent selective pressure favors the emergence of resistance mechanisms (3–6). Through our work, we hope to foster the development of AR LBD targeting therapeutics which prioritize ligand adaptability to enhance long-term efficacy. We anticipate this new approach will yield a class of ligands with distinct potential to improve patient outcomes. We also anticipate that our application of green chemistry will support a rise in sustainable approaches to synthesis within the pharmaceutical industry, making our work to prioritize the approaches in the rational design of CBs increasingly relevant.

## Materials and Methods

### Solid state synthesis methods

Synthesis and ^1^H NMR are described at length in the supplemental methods. Acquisition of crystal structures for the **ClCB** and **BrCB** are as follows: about 10 mg of **ClCB** was dissolved in ethanol and allowed to slowly evaporate. Plate-like crystal suitable for SCXRD was obtained after 24 hr. About 10 mg of **BrCB** was dissolved in benzene and acetonitrile (1:1) and allowed to slowly evaporate. Plate-like crystal suitable for SCXRD was obtained after 48 h. Mercury software 4.2.0 (RRID:SCR_004231) was used to generate representations of crystal structures, stacking, intermolecular interactions, and packing.

### Cell culture reagents and media conditions

CB compounds were prepared as stock solutions in DMSO. The anti-androgens Enzalutamide (MDV3100, Astellas) and Darolutamide (ODM-201, MedChem Express) and solubilized in DMSO (>98% purity). The synthetic androgen R1881 (AbMole) was prepared in DMSO. LNCaP/AR, LNCaP (RRID:CVCL_0395), and MR49F (RRID:CVCL_RW53) cells were maintained in RPMI-1640 10% FBS with the addition of 10uM Enzalutamide for the MR49F cell line. For experimental conditions requiring hormone restriction, LNCaP/AR, LNCaP, and MR49F cells were cultured in RPMI-1640 with 10% CSS FBS. Media with phenol red was used due to weak acidity which aids in solubility of the CBs. All media contained 1xPenStrep, and 0.2% normocin (InvivoGen). For hormone restriction experiments, LNCaP/AR and LNCaP cells were hormone restricted for 24hr prior to treatments, and MR49F cells were cultured in hormone restricted conditions for 72hr prior to treatments. All cell lines tested mycoplasma negative using the MycoStrip Kit (invivogen) throughout experimental use.

### RNA isolation and RT-qPCR

LNCaP/AR, LNCaP, and MR49F cells were seeded into 6-well plates and maintained in hormone restricted conditions. Cells were treated for 24hr prior to collection consistent with previous approaches (20). Total RNA was isolated using the Maxwell 16 RNA purification system (Promega; RRID:SCR_025867). cDNA synthesis was completed using the iScript Reverse transcription supermix for RT-qPCR (BioRad). AR target gene transcription was evaluated by qPCR using validated PrimePCR SYBR primers (BioRad) and SsoAdvanced Universal SYBR Green Supermix (BioRad). Data was normalized to the reference genes TBP and YWHAZ. Data represents the means ± SEM of three independent experiments performed in triplicate.

### Immunocytochemistry and confocal microscopy

Cells were plated into sterilized coverslips coated with poly-D-lysine in 6 well plates and incubated in hormone deprived conditions. Cells were treated for 4hr then fixed and permeabilized with 100% MeOH, blocked in 1% normal horse serum, and stained with an Anti-Androgen Receptor antibody (1:200, abcam227678, RRID:2833098). Secondary antibody incubation was completed with Donkey anti-rabbit Alexafluor 594 (1:200, Invitrogen, A-21207) and cells were counterstained with DAPI (Sigma) for nuclear DNA. Images were captured with a Nikon A1R confocal microscope (RRID:SCR_020317) through a 10µm Z-stack (1.2um step) using NIS-Elements AR software.

Image acquisition settings were kept consistent across treatment groups within cell lines. For nuclear localization studies, 3 images taken at 60x were evaluated in ImageJ for each biological replicate. The JACoP ImageJ Plug-In (RRID:SCR_003070) was used to generate the Mander’s 2 colocalization coefficients using unbiased thresholding through blinded analysis. The nuclear/cytoplasmic fraction was calculated from the proportions of nuclear and cytoplasmic AR. Data represent the means ± SEM of three independent experiments performed in triplicate.

### MTT assay

MTT assays were performed using the Cell proliferation Kit I (Roche). Prostate cancer cells were seeded into wells at 8000 cells/well (LNCaP and MR49F) or 2000 cells/well (LNCaP/AR). CB compounds were prepared in DMSO and applied to cells 24hr after plating. The dosing range for all treatments was 100uM-0.01uM. MTT reagents were added to wells after 96hr of drug treatment. Absorbance was measured using the Varioskan LUX Multi-Mode Microplate Reader (RRID:SCR_26792) at 562nm and corrected by subtracting the absorbance at 650nm. Due to precipitation of CBs at 100uM after addition to media, data points at this concentration were excluded from analysis for all cell lines. Data represents the means ± SEM of three independent experiments performed in triplicate.

### Molecular Docking

Molecular docking studies were conducted using the Molecular Operating Environment (MOE) software from the Chemical Computing group, version MOE2024.06.01 The crystal structure of the wild-type androgen receptor ligand binding domain in complex with S-1 (PDB 2AXA), the T877A mutant with S-1 (PDB 2AX7), and the F877L/T878A mutant with dihydrotestosterone (DHT) (PDB 8FH1) were obtained from the Protein Data Bank (PDB). Prior to docking, all loaded 3D coordinates were corrected. First, polar hydrogens and partial charges were added using the “ Protonate 3D “ function of MOE. For the protonation process, a temperature of 300 K, a concentration of 0.1 mol/L salt in the solvent, and a pH of 7 were specified. Secondly, any missing heavy atoms, alternate geometries, or other crystallographic artifacts were corrected using the “ QuickPrep “ function.

The ligands **ClCB** and **BrCB** were modelized from ChemDraw-generated Simplified Molecular Input Line Entry System (SMILES) and modified using the “ Builder “ tool to define fixed stereochemistry. Both ligands were protonated and energy-minimized using the AMBER10:Extended Hückel Theory (EHT) force field. These structures were included in the input docking database and subsequently docked into previously prepared androgen receptor ligand-binding domains.

The flexible receptor docking was performed by specifying “ Induced Fit “ as the refinement method. The number of placement poses was set to 50, and the number of generated conformations from the refinement stage was set to 5. The charges of all atoms were reassigned under the EHT force field. Final pose scoring was performed based on the London dG free binding energy. For further analysis of the ligand-receptor interactions, the docked poses with the lowest London dG values were chosen.

” Align Sequences and Superpose “ function of MOE allowed representation of the androgen receptor ligand-binding domains from the same angle. “ Measure distances “ function of MOE was used to calculate the interatomic distance between two selected atoms, with the corresponding atoms specified in the manuscript and all distances reported in angstroms (Å).

### Statistics

Ordinary one-way ANOVAs and two-way ANOVAs with Dunnett’s multiple comparisons test, and nonlinear regression were performed using GraphPad/Prism version 10.5.0 (GraphPad Software Inc, RRID:SCR_002798). Graphs in figures 2-4 (2 B-D, 3G, 4B-C) represent sample mean ± SEM n=3. Significance is represented by *p ≤ 0.05, **p ≤ 0.01, ***p ≤ 0.001, and ****p ≤ 0.0001. Nonlin Fit was used to determine the IC50 values in Figure 4A.

## Author Contributions

The manuscript was written through contributions of all authors. A.N.C., C.I.E., L.R.M., and W.A.R. designed research, S.F. and P.L.B. contributed to computational experiments and analysis, A.N.C., C.I.E., and S.F., performed research A.N.C., S.F., and C.I.E. wrote the manuscript. All authors have given approval to the final version of the manuscript.

## Acknowledgments

Research support was provided by the Université de Sherbrooke Center in Green Chemistry and Catalysis (FRQ). We thank the University of Wisconsin-Madison School of Pharmacy Microscopy core for access to the Nikon A1R confocal microscope. The MR49F cell line was provided by Dr. Amina Zoubeidi, and the LNCaP and LNCaP/AR cell lines were provided by Dr. Donald Vander Griend and recommended by Dr. Jordan Vellky. We thank the Ricke Lab and Emily Ricke for laboratory support and pre-submission manuscript review.

## Data Availability Statement

The crystal data that supports the findings of this study are openly available in Cambridge Structural Database at https://www.ccdc.cam.ac.uk, reference numbers 2484979, 2484980, 2484981, and 2484982. All other data are included in the manuscript or SI.

## Supplementary Material

### Supplementary methods

General Synthetic Methods. All reagents were used as purchased. Tetrahydrofuran, 1,4-Dioxane, diethyl ether, Dichloromethane, and Ethyl acetate used in reactions were dried anhydrous. Solvents used for extraction and flash chromatography were reagent grade, purchased from Fisher Scientific. All reactions were carried out under dry N2 atmosphere except where noted. The organic products were extracted with EtOAc and washed with brine dried over Na2SO4 and concentrated under reduced pressure. The product was purified using 10% ethyl acetate in hexanes to yield pure solids. 1HNMR spectra was obtained on 400 MHz Bruker NMR spectrometers.

### Synthesis of (E)-2-Bromo-4-styrylpyrimidine

According to the coupling reaction described by Tan et al (32). 1 trans-2-phenylvinylboronic acid (0.8226 g, 1 eq, 5.6 mmol) was dissolved in 25 mL THF. The catalyst, PdCl2(PPh3)2 (117 mg, 0.03 eq, 0.17 mmol) was added, then potassium phosphate tribasic (2.377 g, 2 eq, 5.6 mmol) was also added. The 2,4-dibromopyrimidine (1.332 g, 1 eq, 5.6 mmol) was dissolved in 5 mL THF and then added to the stirred solution. Water (5 mL) was added. The flask was equipped with a condenser and brought to a reflux for 24 hours. The reaction was quenched with 25 mL water. The organic products were extracted with 3X25 mL EtOAc. They were mixed and washed with brine (30 mL), dried over Na2SO4 and concentrated under reduced pressure. The product was purified using 10% ethyl acetate in hexanes to yield pure 0.8062 g (61% yield) of off-white solid.

1H NMR (400 MHz, DMSO) δ 8.97 (s, 2H), 7.95 (d, J = 16.0 Hz, 1H), 7.80 – 7.73 (m, 2H), 7.49 – 7.32 (m, 4H), 7.26 (d, J = 16.1 Hz, 1H).

The synthesis of (E)-2-chloro-4-styrylpyrimidine and ClCB has been reported as Compound 10 (20).

### Solid state Irradiation of (E)-2-Bromo-4-styrylpyrimidine

Finely ground samples of (E)-2-Bromo-4-styrylpyrimidine (40 mg) were immersed between two Pyrex glass plates and irradiated using a 500 W broadband mercury lamp. 1H NMR spectroscopy was used to monitor the progress of the reaction, and 100% conversion was achieved in 12 h.

1H NMR (400 MHz, DMSO) δ 8.38 (d, J = 5.1 Hz, 2H), 7.37 (d, J = 5.1 Hz, 3H), 7.27 – 7.15 (m, 10H), 7.15 – 7.02 (m, 4H), 4.88 – 4.75 (m, 2H), 4.76 – 4.63 (m, 2H).

## Supplementary Figures

**Supplementary Figure 1.**
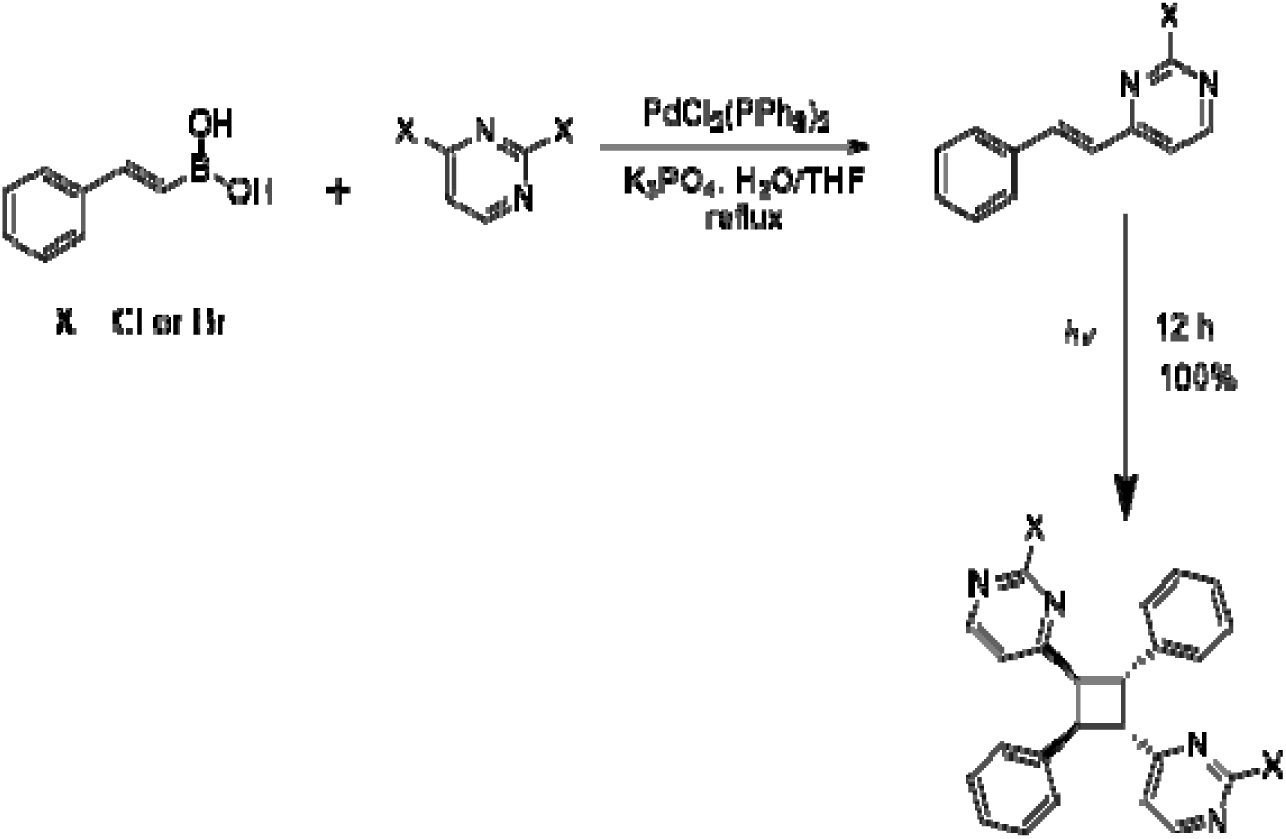
Synthesis scheme for **ClCB** and **BrCB**.

**Supplementary Figure 2.**
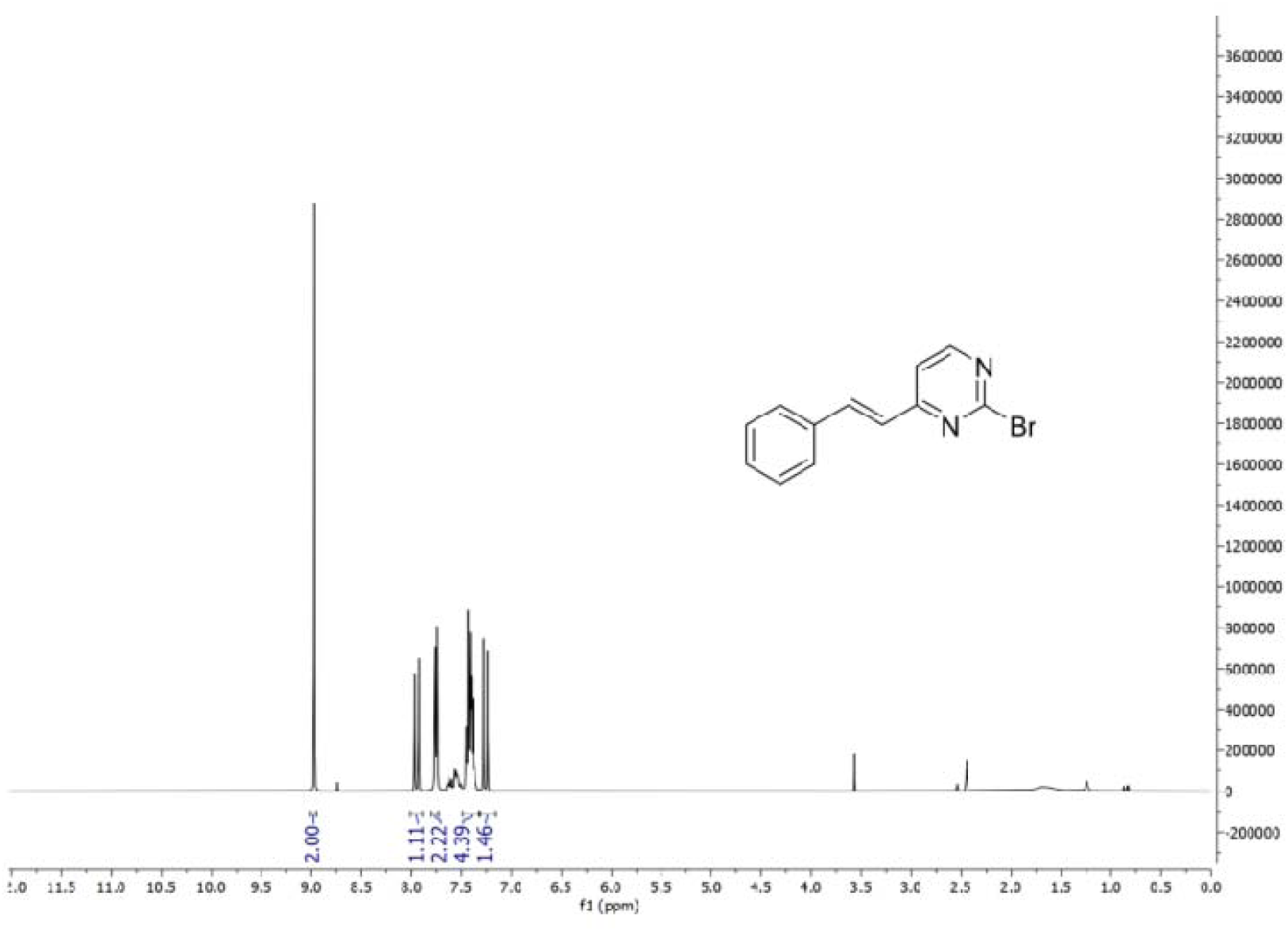
^1^H NMR (400 MHz, DMSO) of (*E*)-2-Bromo-4-styrylpyrimidine.

**Supplementary Figure 3.**
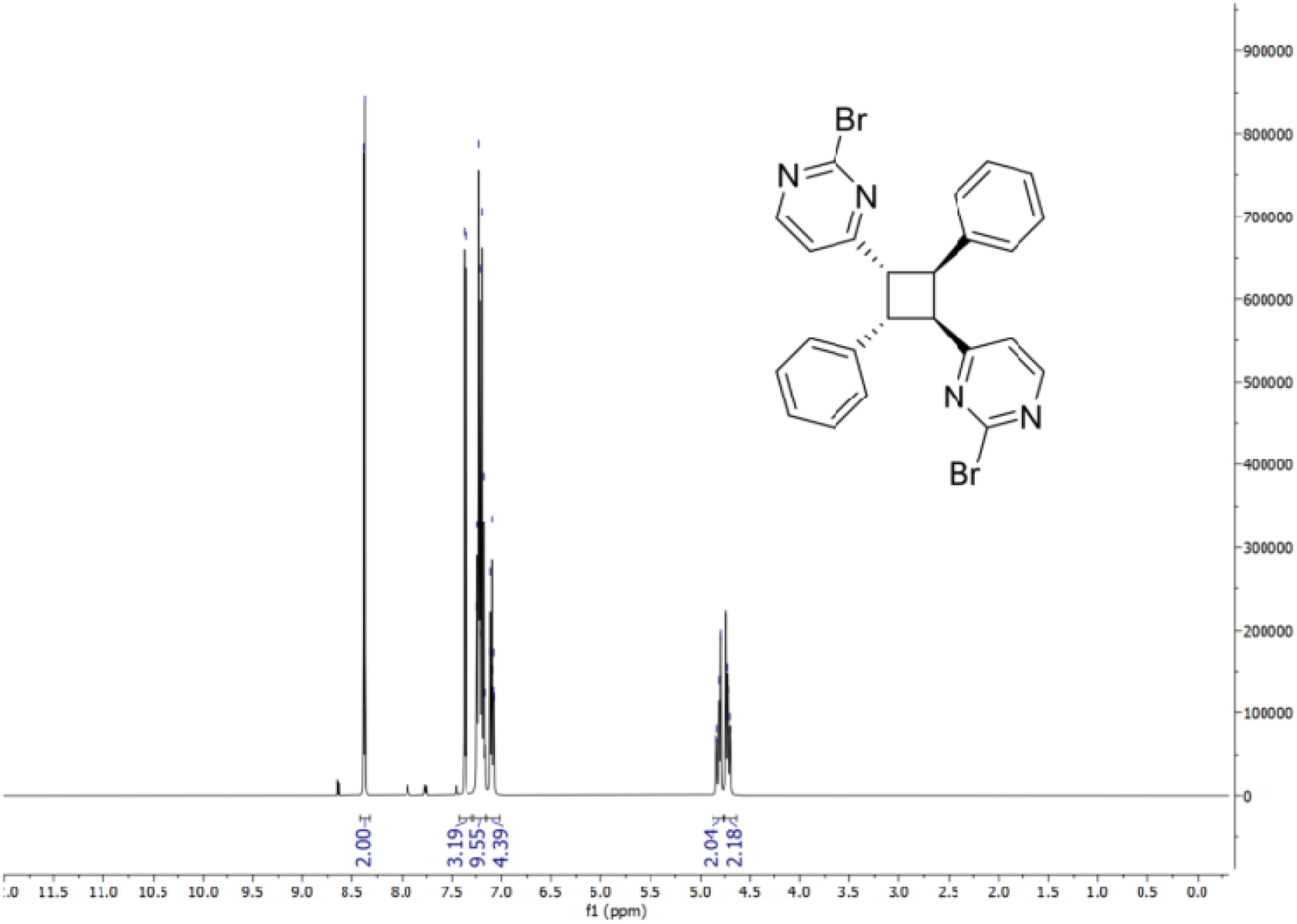
^1^H NMR (400 MHz, DMSO) of **BrCB**.

**Supplementary Figure 4.**
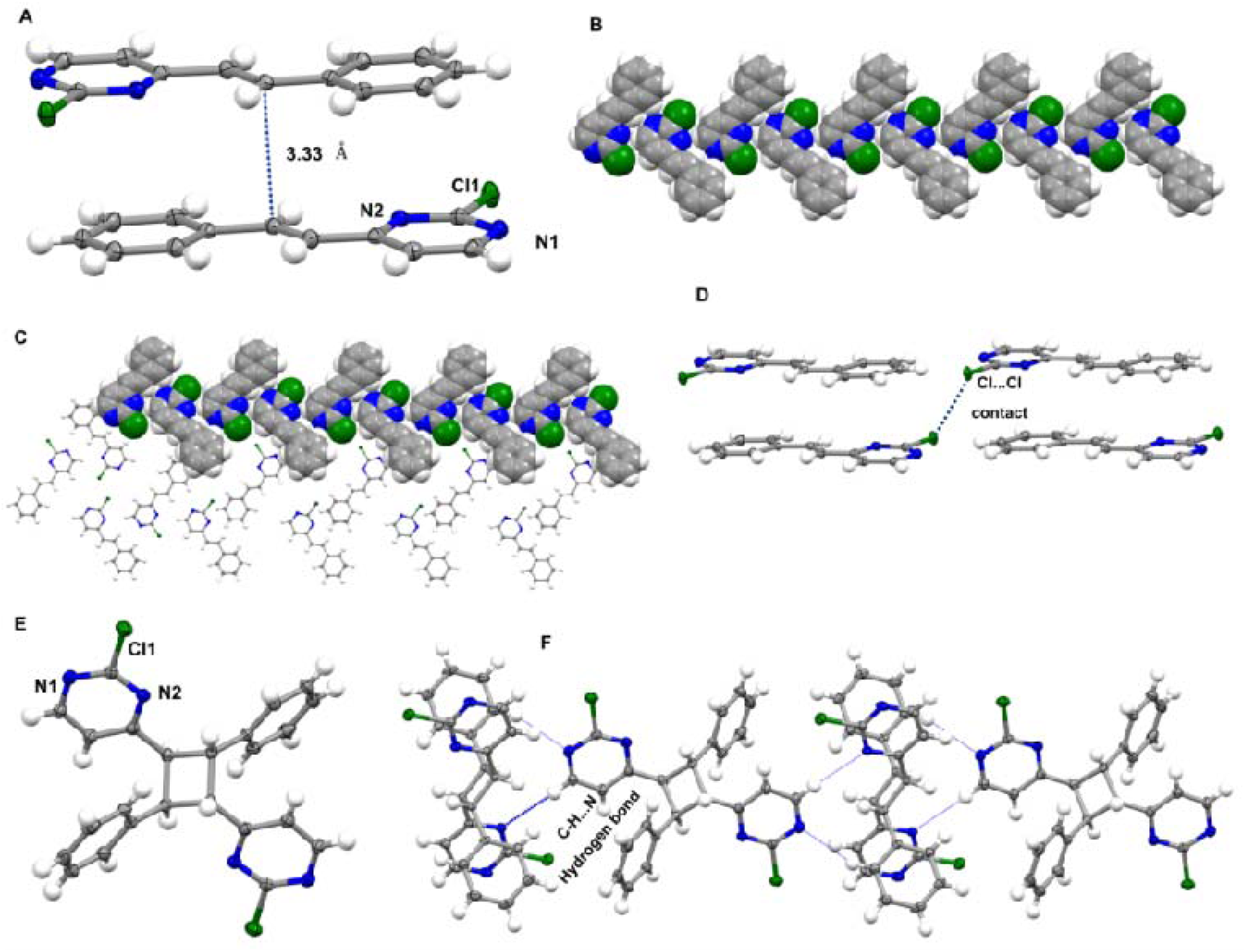
Crystal structures ClSP and ClCB. **(A)** head-to-tail dimer showing the stacking of the C=C bonds of the alkene with 3.33 Å distance **(B)** 1D edge-to-edge view of sheets encompassing neighboring molecules of **ClSP** supported by C-H‧‧‧ N hydrogen bond and **(C)** 2D C-H‧‧‧ N intermolecular interactions **(D)** Cl ‧‧‧ Cl halogen bond of the nearest neighbor **ClSP** (green) **(E)** Single crystal structure for **ClCB** highlighting the *rctt* isomer **(F)** Extended packing of the **ClCB** showing the interaction of the nearest neighbor to form a planar sheet. Corresponding color identifiers for atoms include hydrogen (white), carbon (grey), nitrogen (blue) and chlorine (green). All intermolecular interactions are indicated by dotted blue lines.

**Supplementary Figure 5.**
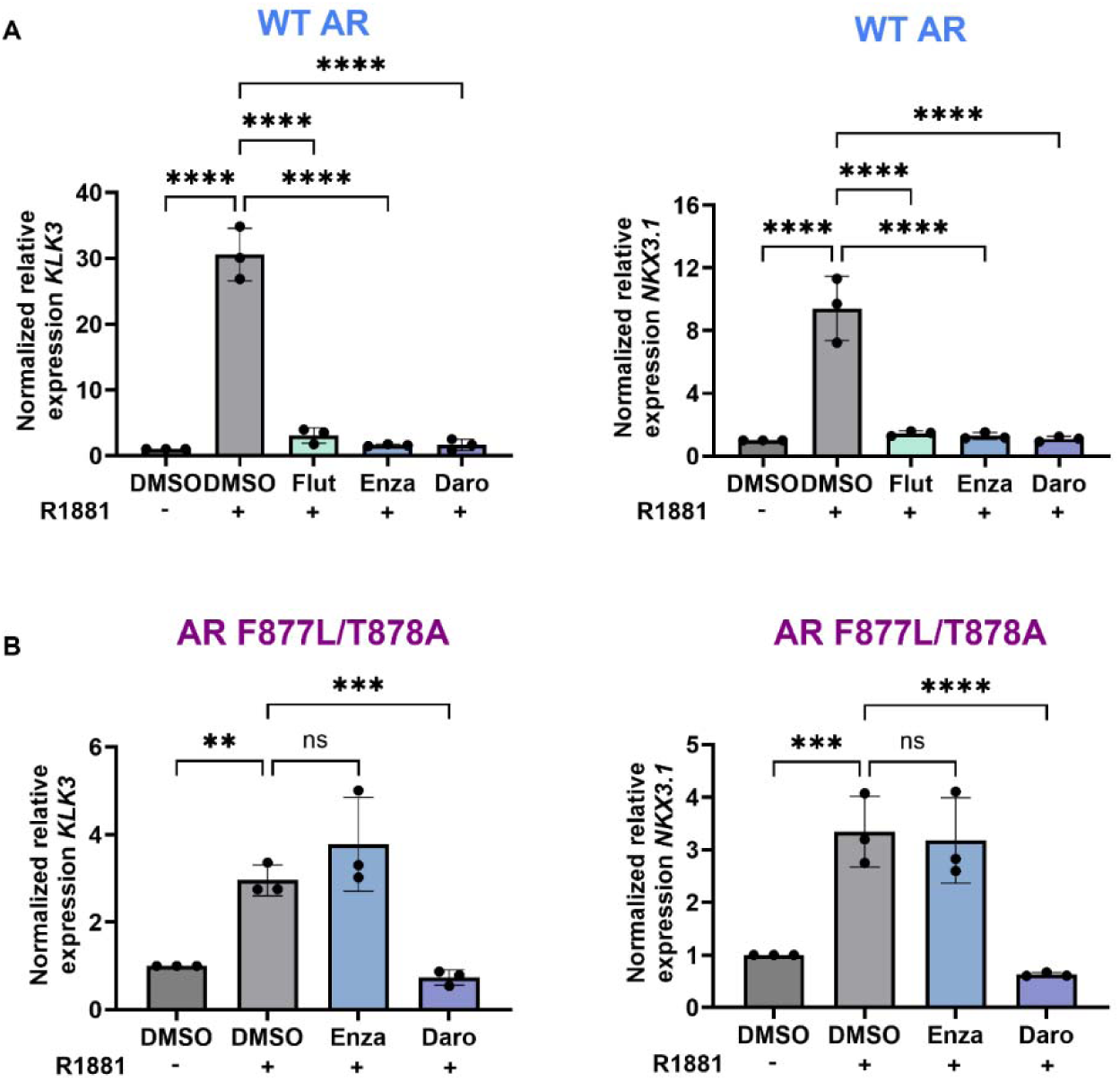
Validation experiments on the transcriptional response of WT AR and AR^F877L/T878A^ to traditional ARSIs. (A) The LNCaP/AR cell line is susceptible to inhibition by flutamide due to codon switch induced to reverse the A878 residue back to T878. qPCR quantification of gene expression in WT AR cells after 24hr exposure to 1nM R1881 or 1nM R1881 + 20µM ARSIs. Notably, flutamide reduces *KLK3* and *NKX3.1* expression with similar ability to second generation anti-androgens. (B) Enzalutamide loses antagonistic function in the MR49F cell line due to the F877L point mutation. qPCR quantification of gene expression in MR49F cells after 24hr exposure to 1nM R1881 or 1nM R1881 + 20µM ARSIs. Enza is unable to block the expression of *KLK3* and *NKX3.1* after exposure to synthetic androgen indicating therapeutic resistance.

**Supplementary Figure 6.**
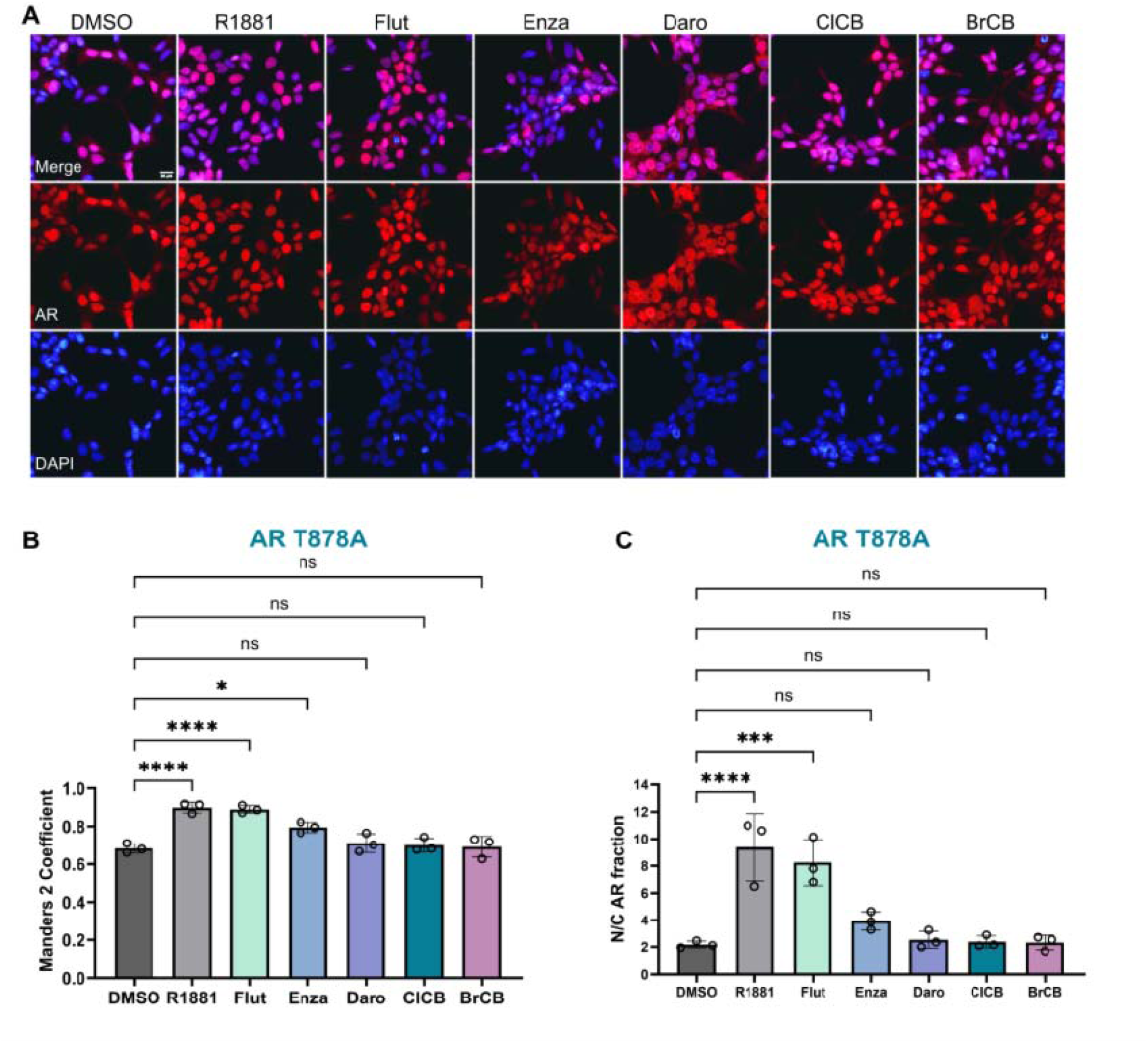
Evaluation of agonistic, partial agonistic, or antagonistic activity of ARSIs and CBs in LNCaP cells. (A) Representative ICC images of AR (red) and changes in protein localization post 4hr treatment with compounds. (B and C) Quantification of AR protein colocalization with the nuclear marker DAPI (blue) by Manders 2 coefficient and representation of the nuclear/cytoplasmic fraction of AR protein. Studies were performed after hormone deprivation to evaluate the presence of agonistic, partial agonistic, or antagonistic activity of each compound in the absence of androgen.

**Supplementary Figure 7.**
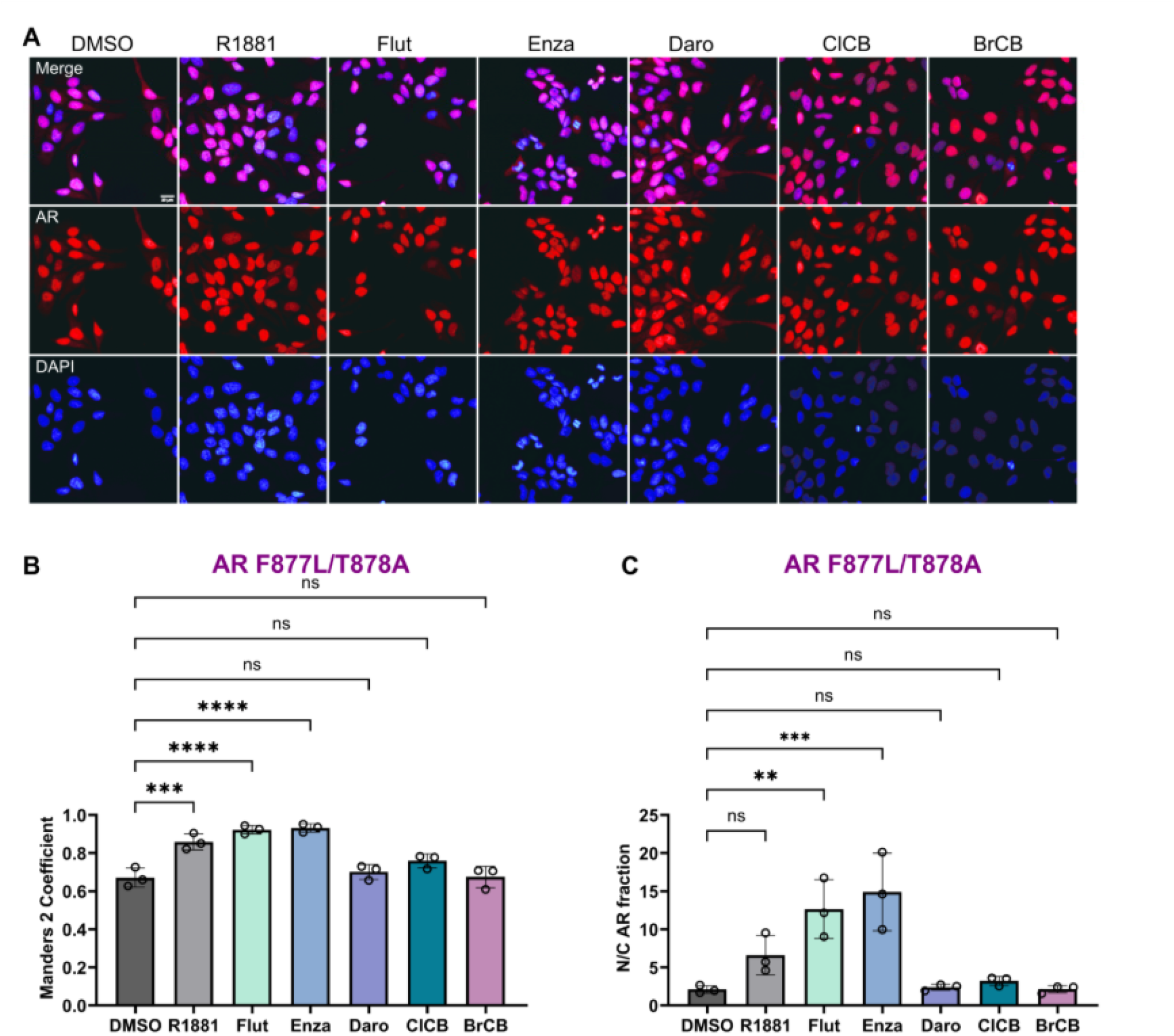
Evaluation of agonistic, partial agonistic, or antagonistic activity of ARSIs and CBs in MR49F cells. (A) Representative ICC images of AR (red) and changes in protein localization post 4hr treatment with compounds. (B and C) Quantification of AR protein colocalization with the nuclear marker DAPI (blue) by Manders 2 coefficient and representation of the nuclear/cytoplasmic fraction of AR protein. Studies were performed after hormone deprivation to evaluate the presence of agonistic, partial agonistic, or antagonistic activity of each compound in the absence of androgen.

**Supplementary Figure 8.**
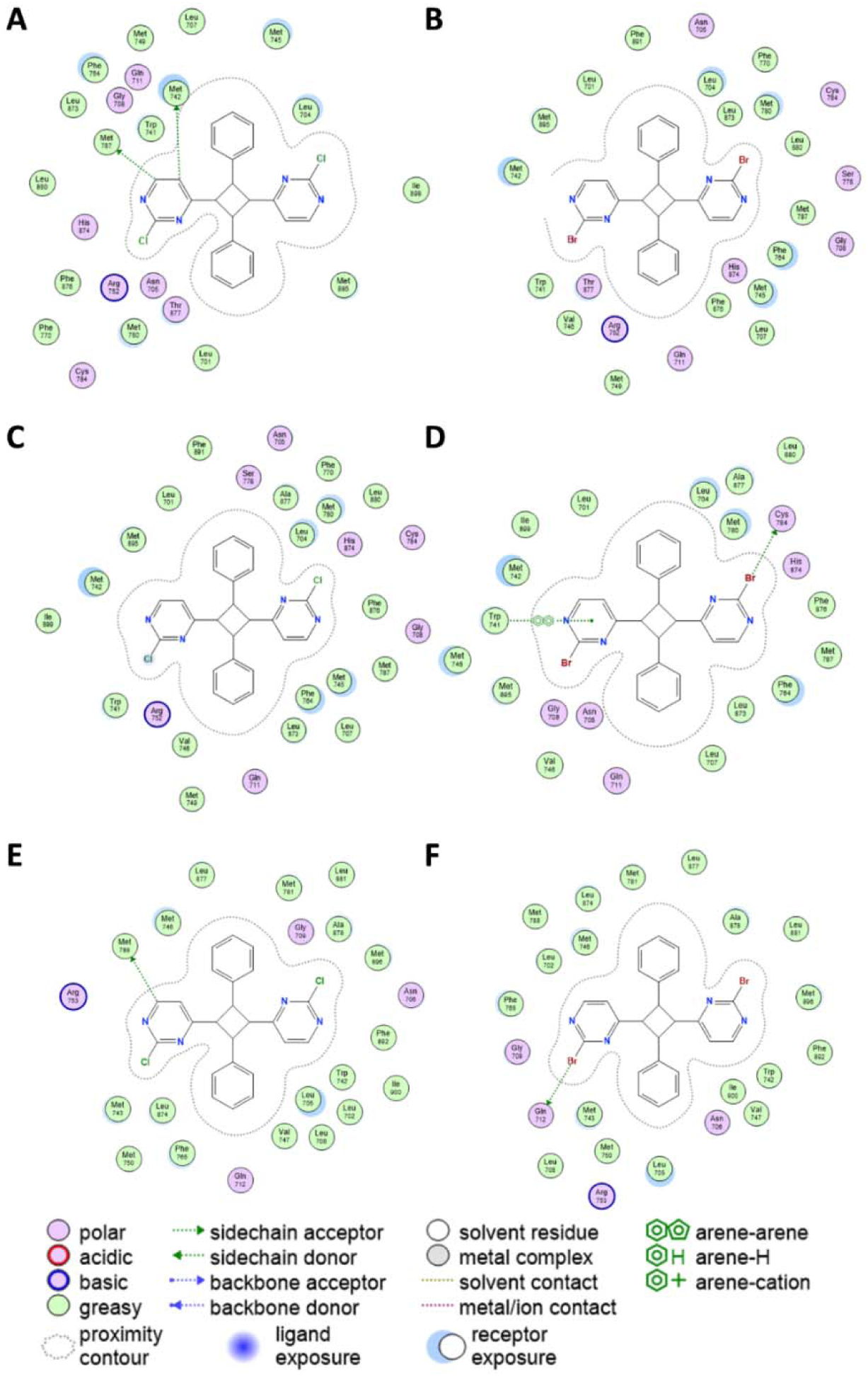
2D interaction diagram of receptor-ligand complex. 2D representation of ClCB (left) and BrCB (right) within the (A, B) WT, (C, D) T877A mutant, and (E, F) F877L/T878A double mutant AR LBD respectively, obtained by docking. The legend interpretation of the ligand-receptor complex interactions is listed at the bottom of the figure. The directional arrows indicate the acceptor/donor nature. The proximity contour represented as a dotted outline that surrounds the ligand, is proportional to the proximity of the ligand to the receptor cavity.

**Supplementary Table 1.**
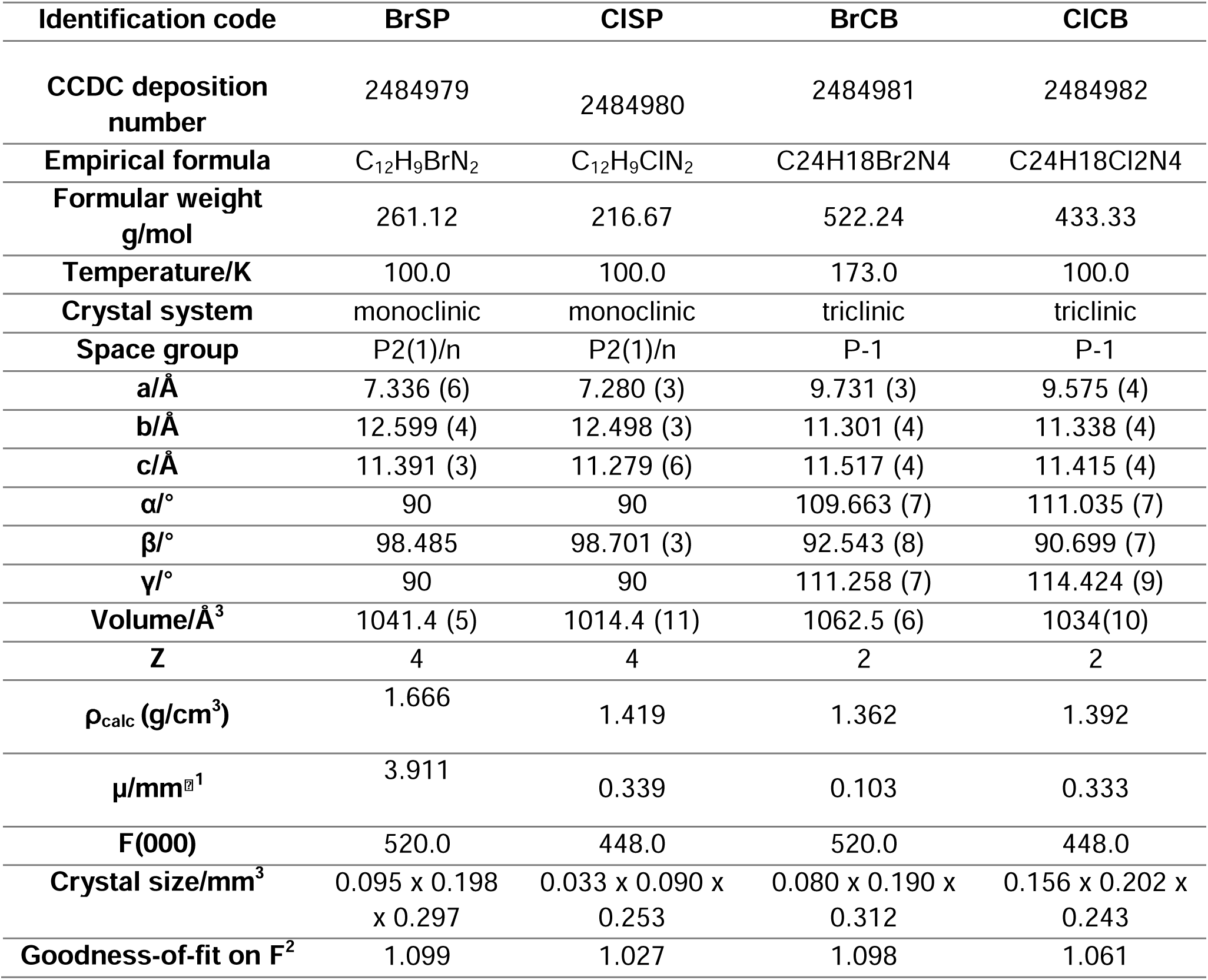
Crystallographic data and structure refinement parameters of ClSP, [(E)-2-Bromo-4-styrylpyrimidine] (BrSP), [(E)-2-Chloro-4-styrylpyrimidine].

**Table S2a.** Interaction report for the ligand **ClCB** in complex with the WT AR LBD.

| WT AR LBD |  | Type of interaction | Distance (Å) | E (kcal/mol) |
| --- | --- | --- | --- | --- |
| Atom | Residue |  |  |  |
| SD | MET742 | H-bond | 3.00 | -0.5 |
| SD | MET787 | H-bond | 2.64 | -0.5 |

**Table S2b.** Interaction report for the ligand **BrCB** in complex with the T877A mutant AR LBD.

| T877A mutant AR LBD |  | Type of interaction | Distance (Å) | E (kcal/mol) |
| --- | --- | --- | --- | --- |
| Atom | Residue |  |  |  |
| - | TRP741 | Aromatic | 4.45 | - |
| SD | CYS784 | H-bond | 3.48 | -0.5 |

**Table S2c.** Interaction report for the ligand **ClCB** in complex with the F877L/T878A double mutant AR LBD.

| F877L/T878A double mutant AR LBD |  | Type of interaction | Distance (Å) | E (kcal/mol) |
| --- | --- | --- | --- | --- |
| Atom | Residue |  |  |  |
| SD | MET788 | H-bond | 2.46 | -0.8 |

**Table S2d.** Interaction report for the ligand **BrCB** in complex with the F877L/T878A double mutant AR LBD.

| F877L/T878A double mutant AR LBD |  | Type of interaction | Distance (Å) | E (kcal/mol) |
| --- | --- | --- | --- | --- |
| Atom | Residue |  |  |  |
| OE1 | GLN712 | Halogen bond | 3.79 | -1.8 |

## References

1. Culig Z, Santer FR. Androgen receptor signaling in prostate cancer. Cancer and Metastasis Reviews. 2014;33(2-3):413–27.

2. Beltran H, Yelensky R, Frampton GM, Park K, Downing SR, Macdonald TY, et al. Targeted Next-generation Sequencing of Advanced Prostate Cancer Identifies Potential Therapeutic Targets and Disease Heterogeneity. European Urology. 2013;63(5):920–6.

3. Steketee K, Timmerman L, Ziel-Van Der Made ACJ, Doesburg P, Brinkmann AO, Trapman J. Broadened ligand responsiveness of androgen receptor mutants obtained by random amino acid substitution of H874 and mutation hot spot T877 in prostate cancer. International Journal of Cancer. 2002;100(3):309–17.

4. Veldscholte J, Ris-Stalpers C, Kuiper GG, Jenster G, Berrevoets C, Claassen E, et al. A mutation in the ligand binding domain of the androgen receptor of human LNCaP cells affects steroid binding characteristics and response to anti-androgens. Biochemical and biophysical research communications. 1990;173(2):534–40.

5. Hara T, Miyazaki J, Araki H, Yamaoka M, Kanzaki N, Kusaka M, et al. Novel mutations of androgen receptor: a possible mechanism of bicalutamide withdrawal syndrome. Cancer Res. 2003;63(1):149–53.

6. Korpal M, Korn JM, Gao X, Rakiec DP, Ruddy DA, Doshi S, et al. An F876L mutation in androgen receptor confers genetic and phenotypic resistance to MDV3100 (enzalutamide). Cancer Discov. 2013;3(9):1030–43.

7. Joseph JD, Lu N, Qian J, Sensintaffar J, Shao G, Brigham D, et al. A clinically relevant androgen receptor mutation confers resistance to second-generation antiandrogens enzalutamide and ARN-509. Cancer Discov. 2013;3(9):1020–9.

8. Gim HJ, Park J, Jung ME, Houk KN. Conformational dynamics of androgen receptors bound to agonists and antagonists. Scientific Reports. 2021;11(1).

9. Borgmann H, Lallous N, Ozistanbullu D, Beraldi E, Paul N, Dalal K, et al. Moving Towards Precision Urologic Oncology: Targeting Enzalutamide-resistant Prostate Cancer and Mutated Forms of the Androgen Receptor Using the Novel Inhibitor Darolutamide (ODM-201). Eur Urol. 2018;73(1):4–8.

10. Ledet EM, Lilly MB, Sonpavde G, Lin E, Nussenzveig RH, Barata PC, et al. Comprehensive Analysis of *AR* Alterations in Circulating Tumor DNA from Patients with Advanced Prostate Cancer. The Oncologist. 2020;25(4):327–33.

11. Liu H, An X, Li S, Wang Y, Li J, Liu H. Interaction mechanism exploration of R-bicalutamide/S-1 with WT/W741L AR using molecular dynamics simulations. Molecular BioSystems. 2015;11(12):3347–54.

12. Liu H, Han R, Li J, Liu H, Zheng L. Molecular mechanism of R-bicalutamide switching from androgen receptor antagonist to agonist induced by amino acid mutations using molecular dynamics simulations and free energy calculation. Journal of Computer-Aided Molecular Design. 2016;30(12):1189–200.

13. Liu H, Wang L, Tian J, Li J, Liu H. Molecular Dynamics Studies on the Enzalutamide Resistance Mechanisms Induced by Androgen Receptor Mutations. Journal of Cellular Biochemistry. 2017;118(9):2792–801.

14. Liu H-L, Zhong H-Y, Song T-Q, Li J-Z. A Molecular Modeling Study of the Hydroxyflutamide Resistance Mechanism Induced by Androgen Receptor Mutations. International Journal of Molecular Sciences. 2017;18(9):1823.

15. Van Der Kolk MR, Janssen MACH, Rutjes FPJT, Blanco-Ania D. Cyclobutanes in Small-Molecule Drug Candidates. ChemMedChem. 2022;17(9).

16. Lee-Ruff E, Mladenova G. Enantiomerically pure cyclobutane derivatives and their use in organic synthesis. Chem Rev. 2003;103(4):1449–83.

17. Xu Y, Conner ML, Brown MK. Cyclobutane and Cyclobutene Synthesis: Catalytic Enantioselective [2+2] Cycloadditions. Angewandte Chemie International Edition. 2015;54(41):11918–28.

18. Carreira EM, Fessard TC. Four-membered ring-containing spirocycles: synthetic strategies and opportunities. Chem Rev. 2014;114(16):8257–322.

19. Sinnwell MA, Groeneman RH, Ingenthron BJ, Li C, MacGillivray LR. Supramolecular construction of a cyclobutane ring system with four different substituents in the solid state. Commun Chem. 2021;4(1):60.

20. Pollock JA, Wardell SE, Parent AA, Stagg DB, Ellison SJ, Alley HM, et al. Inhibiting androgen receptor nuclear entry in castration-resistant prostate cancer. Nature Chemical Biology. 2016;12(10):795–801.

21. Schomaker JM, Delia TJ. Arylation of halogenated pyrimidines via a Suzuki coupling reaction. J Org Chem. 2001;66(21):7125–8.

22. Abate-Shen C, Nunes De Almeida F. Establishment of the LNCaP Cell Line – The Dawn of an Era for Prostate Cancer Research. Cancer Research. 2022;82(9):1689–91.

23. Horoszewicz JS, Leong SS, Chu TM, Wajsman ZL, Friedman M, Papsidero L, et al. The LNCaP cell line--a new model for studies on human prostatic carcinoma. Prog Clin Biol Res. 1980;37:115–32.

24. Coleman DJ, Van Hook K, King CJ, Schwartzman J, Lisac R, Urrutia J, et al. Cellular androgen content influences enzalutamide agonism of F877L mutant androgen receptor. Oncotarget. 2016;7(26):40690–703.

25. Tran C, Ouk S, Clegg NJ, Chen Y, Watson PA, Arora V, et al. Development of a Second-Generation Antiandrogen for Treatment of Advanced Prostate Cancer. Science. 2009;324(5928):787–90.

26. Chen Y, Clegg NJ, Scher HI. Anti-androgens and androgen-depleting therapies in prostate cancer: new agents for an established target. The Lancet Oncology. 2009;10(10):981–91.

27. Nangia AK, Desiraju GR. Crystal Engineering: An Outlook for the Future. Angewandte Chemie International Edition. 2019;58(13):4100–7.

28. Freund R, Lachelt U, Gruber T, Ruhle B, Wuttke S. Multifunctional Efficiency: Extending the Concept of Atom Economy to Functional Nanomaterials. ACS Nano. 2018;12(3):2094–105.

29. Bolla G, Sarma B, Nangia AK. Crystal Engineering of Pharmaceutical Cocrystals in the Discovery and Development of Improved Drugs. Chem Rev. 2022;122(13):11514–603.

30. Crawford ED, Schellhammer PF, McLeod DG, Moul JW, Higano CS, Shore N, et al. Androgen Receptor Targeted Treatments of Prostate Cancer: 35 Years of Progress with Antiandrogens. J Urol. 2018;200(5):956–66.

31. Hussain M, Saad F, Sternberg CN. Enzalutamide in Castration-Resistant Prostate Cancer. N Engl J Med. 2018;379(14):1381.

32. Tan JQ, Chang JH, Deng MZ. The facile route to stereodefined alkenyl-substituted pyrimidines. Synthetic Commun. 2004;34(20):3773–83.

